# Calcium dynamics tune timing of developmental transitions to generate evolutionarily divergent axon tract lengths

**DOI:** 10.1101/2024.12.28.630576

**Authors:** Feline W. Lindhout, Hanna M. Szafranska, Luca Gulglielmi, Ivan Imaz-Rosshandler, Jerome Boulanger, Maryam Moarefian, Kateryna Voitiuk, Natalia K. Zernicka-Glover, Ulrike Schulze, John Minnick, Daniel J. Lloyd-Davies Sánchez, Ilaria Chiaradia, Laura Pellegrini, Mircea Teodorescu, Madeline A. Lancaster

## Abstract

The human brain has undergone an evolutionary expansion in size, both in terms of cell numbers and the size of cellular structures, including axon tracts. Human brain development also progresses slowly and takes particularly long. However, the functional relevance of slowed timing, and whether it is responsible for these changes in size, remains unknown.

Here, we investigate this by studying axon tract development in human and mouse brain organoids. We demonstrate that human axon tracts grow ∼2x more slowly than those of mice, reflecting their slowed tempo, but that this actually leads to shorter human axons, not longer. To overcome the effect of slowed tempo, human axons have a more prolonged growth duration that enables them to project farther despite their slower growth rate. Using a combination of single-cell RNA sequencing and live imaging, we demonstrate that the prolonged duration involves a different mechanism to that controlling tempo and is driven by calcium dynamics. Human axons exhibit a reduced calcium influx compared to mouse, mediated by L-Type voltage-gated calcium channels. Stimulating this calcium influx in human neurons triggers earlier cessation of growth, leading to shorter axon tracts similar to those of mouse. We further show that increasing calcium speeds up the transition to the synaptogenesis stage. Thus, calcium regulation sets the timing of transitions to disproportionately extend developmental duration, thereby enabling evolutionary expansion of human neurons.

## INTRODUCTION

Many features of the human brain are distinguished by scaling differences of conserved structures. Our brain is larger than that of other primates and rodents, composed of more cells, and homologous cell types show expanded morphologies ^1–18^. This includes cortical neurons with increased dendritic arborizations, synapse numbers, and longer axon tracts, together fostering enhanced connectivity (Fig 1a) ^9,10,14–18^. These scaling differences are particularly pronounced in regions that have undergone functional specialisation in the human brain. For example, the prefrontal cortex shows a disproportionate increase in size, white matter, and connectivity compared with primary cortices, a feature not observed in monkeys or rodents ^9,10,14–17,19–22^. Thus, identifying the mechanisms driving species-specific scaling differences during brain development is central to understanding human brain evolution.

**Figure 1:**
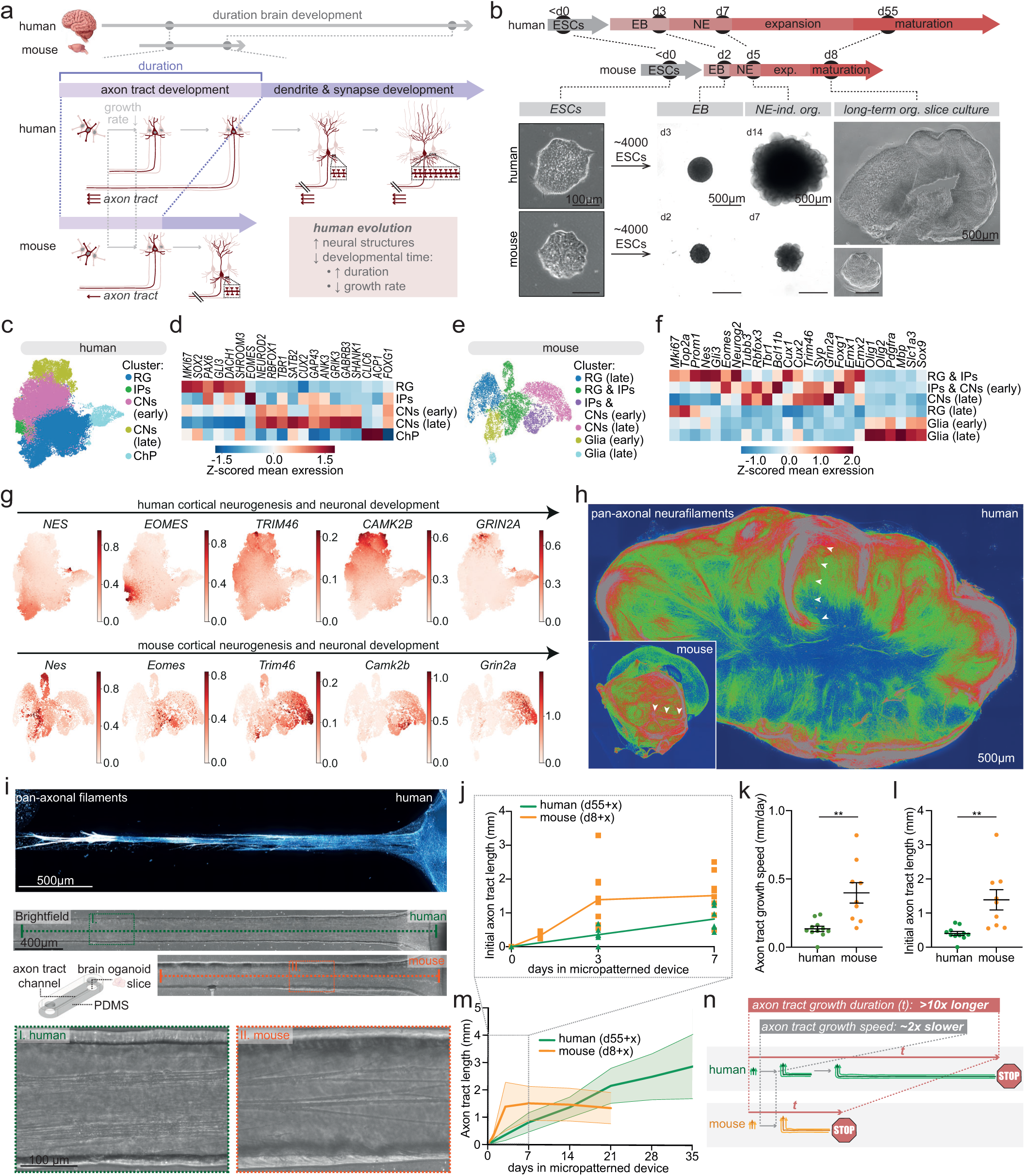
Human brain organoids exhibit longer axon tracts and increased axon tract development duration compared to mouse independent of environmental context. a. Schematic illustration of a hypothesised model where prolonged developmental duration, despite slower growth rates, may explain the expanded neuronal morphologies in human evolution. b. Protocol timeline for the generation of unpatterned long-term human and mouse brain organoid slice cultures using the same protocol steps for both species, with the timing of each step tailored to the species-specific timing ^47^. Brightfield images reflect representative images at different steps of the protocol. c. UMAP embedding of the human brain organoid dataset covering 13 timepoints between days 57-93. d. Heatmap of representative marker genes of clusters in (c). e. UMAP embedding of the mouse brain organoid dataset, subclustered with cortical neurogenesis and gliagenesis clusters in (Extended Data Fig 1i), covering 8 timepoints between days 10-22. f. Heatmap of representative marker genes of clusters in (e). g. Feature plots of SOX2 (neural progenitors), EOMES (intermediate progenitors), NFASC (axon initial segment), CAMK2B (synapses) and GRIN2A (mature synapses) in human and mouse datasets. h. Long-range axonal tracts with similar morphologies in a day 69 human and a day 15 mouse ALI-CO immunostained for pan-axonal neurofilaments, with axon tracts extending further in humans (indicated by arrowheads). i. Top: human organoid slice culture placed in a micropatterned device at day 55, fixed 35 later, respectively, and immunostained for pan-axonal filaments. Bottom: schematic illustration of micropatterned PDMS devices with long channels enabling analysis of outgrowing axon tracts from brain organoid slices. Brightfield images of human (day 55+35) and mouse (day 8+8) organoid slice cultures with an axon tract extending into the channel of the micropatterned device, with zooms showing self-organised axon bundling. j. Zoom of timepoints day 0-7 of plot in (m). m. Quantifications of initial axon tract length after 3 days of human and mouse brain organoid slice cultures in micropatterned devices. Hm: n=11; ms: n=9. See (m) for statistic details. k. Quantifications of initial axon tract outgrowth rates (days 0-3) of human and mouse brain organoid slice cultures in micropatterned devices over time. Hm: day n=10; ms: n=9. l. Quantifications of axon tract length over time of human and mouse brain organoid slice cultures in micropatterned devices. Hm: n=7; ms: n=9 (d0,3,7,14), n=4 (d1,21). Mixed-effects model P<0.0001 for time, p=0.1991 for species (average length across timepoints). Post-hoc Šidák Multiple comparison test for following timepoints; adjusted P value d3: 0.0031 (shown in m), d7: 0.0655, 14: 0.9844. l. Schematic illustration summarising findings. ESC, embryonic stem cells. EB, embryonic body. NE, neuroepithelium. Exp., expansion. Ind., induced. Org., organoid. RG, radial glia. IPs, intermediate progenitors. CNs, cortical neurons. ChP, choroid plexus. Data shown as mean ± SEM (k,l) or mean ± SD (m); \*\**P* < 0.01 as determined by unpaired *t* tests unless stated differently.

Whilst the mechanisms driving the expansion of cell numbers and their morphologies in the human brain remain unclear, there is a consistent correlation with the disproportionately prolonged timeline of human brain development. Across many species, brain size consistently correlates with gestation time ^23^. Moreover, humans have evolved an additional extension of brain maturation, the process coinciding with neuron development, which is disproportionately longer compared with other primates and rodents even when corrected for lifespan differences^23–31^. Together, this raises the hypothesis that the expansion of the human brain and its underlying neurons may result from slowed developmental time, or bradychrony ^23^. Species-specific developmental timing is highly robust, as human neurons retain their slow maturation pace when differentiated *in vitro* and even when transplanted into rodent brains ^32–34^, suggesting that whatever the mechanism is, it is also cell-intrinsic.

Efforts to uncover mechanisms controlling timing have largely focused on tempo, which relates to the rate at which developmental processes occur. These studies have revealed a consistent ∼2x slower rate in human cells than in mice, driven by species-specific differences in protein turnover and metabolism ^35–38^. Along these lines, the slower maturation rate seen in human neurons has been shown to relate to differences in mitochondria metabolism ^39^ and epigenetics ^40^. However, a slower rate alone does not explain evolutionary expansion as it would lead to the opposite developmental outcome, with reduced proliferation and growth rates resulting in fewer cells with less elaborate morphological structures (Fig 1a, Extended Data Fig 1a). Indeed, in differentiating human neurons, perturbing metabolism or epigenetics to accelerate maturation as in the mouse led to outcomes with larger morphological structures ^39,40^, rather than smaller, a result that does not align with the evolutionary pattern.

Instead, a temporal feature that could account for the scaling differences of the human brain is duration. Humans have evolved a much more prolonged duration of brain development than can be explained by the ∼2-fold slower tempo, taking several decades to reach full maturity ^23–31^. This prolonged duration is observed at each step of neurodevelopment, including the process of neuronal morphogenesis,starting with neurite outgrowth and axonogenesis, followed by dendritogenesis and synaptogenesis (Extended Data Fig 1b) ^24–30,41–45^. The extended duration of these processes may enable neurons to form relatively more of these structures despite a slower rate of their growth, by spending an even longer time in each developmental window (Fig 1a, Extended Data Fig 1a). Thus, understanding the scaling differences of the human brain, including morphological expansion of neurons, may require elucidating mechanisms underlying extended developmental duration.

Investigating mechanisms underlying duration is challenging in existing human models precisely because of the extended time of human neuron development, taking many years. However, axon tract development is a process of neuron morphogenesis completed during embryogenesis, making it a more accessible process to study. The expansion of axon tracts is also a perfect example of the evolutionary elaboration of neuronal structures, as they have expanded relatively more compared to other neuronal domains ^18,22,46^. However, whilst other maturation hallmarks are easier to read out using typical human neural *in vitro* models, it is considerably more challenging to analyse axon lengths with these systems. Here, we used modified human and mouse brain organoid slice cultures that robustly form self-organised axon tracts and combined these with tailored micropatterned devices to mechanistically explore the link between species-specific axon tract morphology, developmental duration and growth rates. Using these methods, we found that whilst axon outgrowth rates in humans are ∼2x slower compared to mice, the duration of axon tract elongation is disproportionally longer, ultimately leading to increased axon tract lengths. In human organoids, developing neurons consistently exhibit a lower calcium influx, mediated by L-Type voltage-gated calcium channels (VGCC), compared to similar stages in mice.

Increasing the L-Type VGCC-mediated calcium influx in human neurons mimicked the mouse phenotype and led to a shortening of the axonogenesis time window, as well as shortened axon tracts, due to an earlier transition from axon outgrowth to synaptogenesis. Thus, this work has uncovered species-specific differences in calcium dynamics, which drive the formation of evolutionarily divergent axon tract morphologies by tuning the timing of cell state transitions and with it, developmental durations.

## RESULTS

### Stage-matching cortical neurodevelopment in unguided human and mouse brain organoids by morphometric and transcriptomic profiling

To investigate evolutionary timing differences in axon tract outgrowth and the eventual morphology of axon tracts, we used human and mouse brain organoids generated using a previously described method ^47,48^ as these closely resemble *in vivo*-like developmental duration, without extrinsically driving developmental progression by small molecules that seem to promote earlier maturation ^49^. In order to generate comparable organoids of the proper cerebral cortical identity, human and mouse brain organoids were generated using identical media, with the only difference being the timing of media changes to match the intrinsic species-specific timing and ensure that organoids reached comparable neurodevelopmental stages with proper brain identity and tissue morphology (Fig 1b) ^47^.

Over the course of their development, the organoids exhibited a gradual size divergence, which, despite their overall small size compared with the *in vivo* brain, culminated in a substantial difference in size between the human organoids and the mouse (Fig 1b).

To study axon tract development, we generated human and mouse long-term air-liquid interface cerebral organoid (ALI-CO) cultures, which we have previously shown to successfully recapitulate these relatively later stages of neuronal development with improved cell survival ^47,48^. To properly compare axon tract development between the two species, we sought to align the temporal progression of human and mouse organoids so that the initial timepoint in each would be the onset of axon development. Axon development begins when neurons undergo a symmetry break, transitioning from an unpolarized to a polarized state, involving the accelerated outgrowth of only the future axon. This is the first hallmark of a developing axon ^50^, with other axon-specific features such as axon initial segment assembly appearing ∼1-2 weeks later in human neurons ^51^. We identified this transition using a sparse viral labelling approach enabling evaluation of individual morphologies of the different neuronal polarity states in organoids (Extended Data Fig 1c,d). In both human and mouse organoids, we detected unpolarised neurons with similar neurite lengths, and polarised neurons with axon lengths exceeding those of other neurites (Extended Data Fig 1d). Similar to *in vivo*, neuron development in brain organoids is asynchronous. Thus, we aimed to identify the timepoint where mouse and human exhibited similar proportions of polarized and unpolarized neurons. In human organoids sliced and transduced with a GFP+ virus at day 55 and analysed two days later, approximately half of the neurons were still unpolarized, and an equivalent proportion was observed in mouse organoids sliced at day 8 (Extended Data Fig 1e). At these matched timepoints, the total axon population was also at a comparable developmental stage, with numerous axons present but still individually distinguishable as single projections (Extended Data Fig 1f).

To further validate both identity and stage, we performed single-cell RNA sequencing of 13 human and 8 mouse timepoints, capturing developmental progression from the identified matched stages and onwards (Extended Data Fig 1g,h). Both human and mouse brain organoids displayed forebrain identities, exhibiting a wide range of neural cell types and complexity, as expected ^48,52,53^ (Fig 1c,d, Extended Data Fig 1g-i). Cluster analysis revealed cell populations expressing markers of cortical progenitors and neurons in both human and mouse, including radial glia, intermediate progenitors, deep layer and upper layer neurons (Fig 1c,d, Extended Data Fig 1h,i). Additionally, in human organoids we identified a distinct cluster representing the choroid plexus, which also forms within the forebrain, similar to our previous reports (Fig 1c,d) ^48^. In mouse organoids, where we also captured relatively later developmental stages, we already observed a cluster containing developing astrocytes and oligodendrocytes, which, as expected, predominantly originated from older samples (Extended Data Fig 1h,i).

In mouse organoids, we observed additional clusters matching derivatives of the diencephalon, a subdivision of the forebrain, consistent with previous reports ^54^, which in our dataset included retinal pigmented cells, retinal photoreceptors and retinal inhibitory neurons (Extended Data Fig 1h,i). Importantly, while cortical neurons were abundantly present at the stage-matched timepoints in both human and mouse organoids, retinal neurons in the mouse organoids emerged later, thus the stage-matched timepoints for axon outgrowth capture cortical neuron development (Extended Data Fig 1j). Given our primary interest in comparing cortical neuronal development, we subsetted the mouse dataset with the telencephalic clusters, encompassing neural progenitors, cortical neurons and glia, for further analysis (Fig 1e,f).

In addition to distinct cell types, we also identified cortical neurons at early and late developmental stages, corresponding to axonogenesis and synaptogenesis respectively, in both human and mouse datasets (Fig 1c-f). Feature plots of neurodevelopmental markers revealed the expression of genes involved in axon outgrowth and axon initial segment assembly during the transition from neural progenitors to neurons, followed by a subsequent upregulation of synaptic genes (Fig 1g). In a small subset of the most mature neurons in both the human and mouse datasets, we could already detect the mature synapse marker GRIN2A (Fig 1g), together indicating a broad coverage of the neuronal developmental stages from axonogenesis to synaptogenesis in these datasets. Thus, we successfully captured and stage-matched cortical neuron development in mouse and human organoids, based on morphometric and transcriptomic profiling, to allow accurate comparison of developmental timing and morphological outcome.

### Human organoid axon tracts grow longer than mouse

We next investigated whether human and mouse organoids exhibit species-specific differences in axon tract development, from the initial onset through subsequent developmental stages. We found that newborn human and mouse neurons, when still at the start of neuron development, initially displayed similar morphologies, including comparable neurite lengths and soma areas (Extended Data Fig 1d,k-o). Of note, axons, but not other neurites, already showed a trend indicating a modest difference in growth between human and mouse neurons, with more growth in mouse within one day, indicating a seemingly accelerated growth compared to human (Extended Data Fig 1k). Moreover, similar to previous reports, human and mouse neurons showed a subtle trend toward a decreased soma area upon establishing neuronal polarity or in timepoints corresponding with the onset of neuronal polarity (Extended Data Fig 1n,o)^51^. Once neuronal polarity was established, and the ALI-COs were allowed to reach a more mature stage, organoids developed self-organised long-range axon tracts projecting to distant regions within the ALI-CO in both human and mouse, as described previously (Fig 1h) ^47,48^. A notable difference was that human axon tracts took longer to appear and seemed to have extended farther than those of mice, reminiscent of *in vivo* differences in axon tract length and timing.

In the brain, distinct axon trajectories are established due to local tissue variations, such as differences in stiffness or chemoattractant and chemorepellent levels, that influence growth cone turning events and outgrowth rates ^55,56^. Across species with different brain sizes, homologous axon trajectories can be identified and follow similar paths with the main difference being their length. Such a difference could be due to cell intrinsic differences of axon growth as well as species-specific factors in the tissue environment. To rule out these extrinsic factors and identify intrinsic regulators of timing, we sought to establish a method to examine isolated outgrowing axon tracts from each species grown in an identical and controlled environment. We engineered micropatterned devices from molds generated using photolithography, with chambers for brain organoid slices connected to a 5 mm long channel, to capture outgrowing axon tracts and allow for their unrestricted growth (Fig 1i, Extended Data Fig 2a-c). Within a few days after placing a human brain organoid in one of the chambers, individual axons started to grow out into the channel and continued to extend over subsequent weeks whilst forming large bundles, similar to those observed in ALI-COs (Fig 1h,i). The tract was almost completely composed of axons with a few cells migrating into the tract once axons started to bundle ^57,58^ (Extended Data Fig 2d,e). We observed a similar developmental axon tract pattern of mouse brain organoids in micropatterned devices, but with the key difference that axon tracts from mouse organoids did not extend as far compared to human (Fig 1i).

### Species-specific differences in axon growth duration, not rate, explain differences in length

To identify how the species difference in length comes about, we examined axon tract development over time and observed that the longer axon tracts of human organoids were not due to increased axon outgrowth rates (Fig 1j). In fact, axon outgrowth rates were 2x slower in human organoids (Fig 1j,k), consistent with the trend observed in ALI-COs (Extended Data Fig 1k), resulting in initially shorter instead of longer axon tracts in human organoids (Fig 1l). However, the duration of axon tract growth was disproportionately extended in human compared to mouse, and exhibited a much greater difference in duration than growth rate (Fig 1m), with axon tract growth starting to plateau after about 3 days in mouse compared to axon tracts still extending after about 5 weeks in human, allowing human axon tracts to surpass those of mouse and extend substantially farther (Fig 1j,k).

While micropatterned devices represent an *in vitro* context distinct from *in vivo* conditions, applying the same model system to both species enabled us to examine human and mouse axons in an internally controlled manner with species as the remaining variable.

Reassuringly, while reports on axon growth of primary neurons are limited for humans, the axon elongation window that we observed falls within the 2 days to ∼1-2 weeks range reported for primary rodent neurons, yielding axon lengths of 0.3–1.5 mm, across *in vivo* and *in vitro* studies ^59–63^. By capturing species-specific differences in axon tract developmental timing and morphology, these data further support the notion that this process is governed by cell-intrinsic mechanisms. Together, these data indicate that different aspects of developmental time have not slowed uniformly in human, but rather developmental duration has extended disproportionately through cell-intrinsic mechanisms, providing an explanation for the expansion in axon tract length despite slowed growth rates (Fig 1n). Thus, understanding evolutionarily divergent size differences requires elucidating the mechanisms driving the extension of developmental duration.

The finding that species-specific axon tract length is driven by cell-intrinsic mechanisms suggests that axons may have an inherent growth time window, during which they are not yet receptive to environmental interactions that may direct the transition from axon outgrowth to synapse formation. This also explains how, in ALI-COs as well as *in vivo*, axons initially pass by neurons of similar identities to those they will eventually target. To test this, we examined the effect on the duration of axon outgrowth when presented with a target tissue whilst still in the axon extension stage. Using similar micropatterned devices, but with shorter channels, we placed a human ALI-CO expressing mCherry in one chamber, and a non-fluorescently labelled human ALI-CO in the other chamber (Extended Data Fig 2f). After ∼12 days in culture, corresponding with early axon tract elongation stages for human organoids (Fig 1m), axons had reached the target tissue, but this did not result in axon outgrowth halting. Instead, axons grew over the target tissue and continued to do so over subsequent days (Extended Data Fig 2f). This was also observed when using mouse spinal cord explant as target tissue, which we have previously shown to be functionally innervated by outgrowing axon tracts from human ALI-COs ^48^, with axons from human brain organoids labelled with Marcks-GFP extending over the mouse spinal cord tissue whilst in their axon outgrowth stage (Extended Data Fig 2g). Altogether, these data indicate that developmental duration consistently correlates with species-specific differences in axon tract length, irrespective of environment and target tissue interactions, further supporting that differences in developmental duration underlie species-specific axon tract lengths through cell-intrinsic mechanisms.

### Pseudotime analysis reveals distinct expression profiles of neuronal calcium channels in human

The duration of axon development is determined by the timing of the cell state transition from axonogenesis through to synaptogenesis. To identify regulators influencing the timing of this and potentially other neurodevelopmental transitions, we performed pseudotime analysis of the scRNA-seq datasets to detect genes expressed throughout neurodevelopment and already prior to synaptogenesis. In the human organoid dataset, this analysis resolved two main trajectories: cortical neurogenesis and choroid plexus development (Fig 1c, Fig 2a).

**Figure 2:**
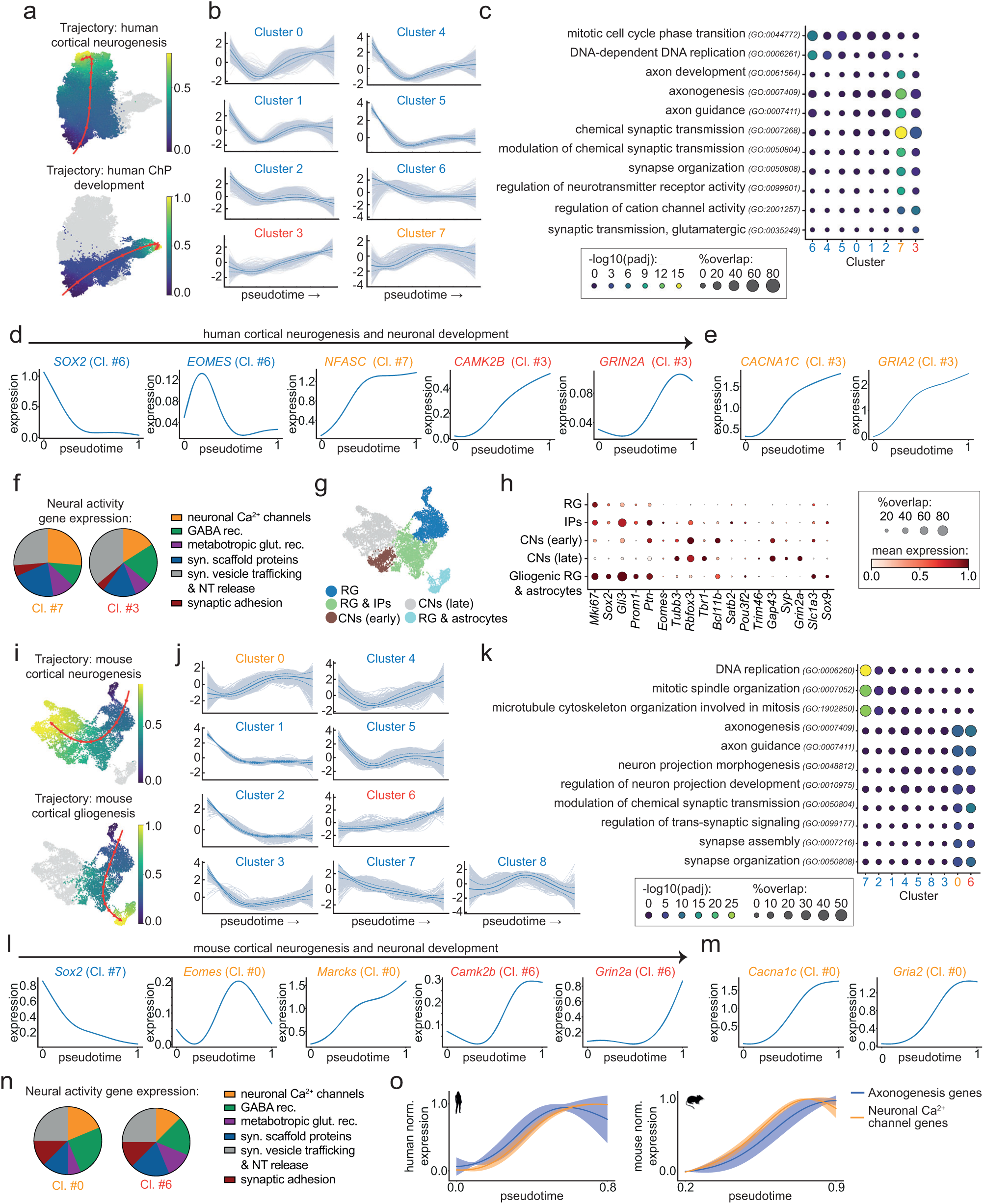
A subset of calcium-permeable ion channels show increased expression during early neuron development in human and mouse. a. Pseudotime trajectories identified in the human RNA-seq dataset. b. Gene clustering based on differential expression patterns along the human cortical neurogenesis pseudotime trajectory. c. Dotplot of GO term enrichment analysis of gene clusters in (b). d. Human pseudotemporal expression patterns of SOX2 (neural progenitors), EOMES (intermediate progenitors), NFASC (axon initial segment), CAMK2B (synapses) and GRIN2A (mature synapses). e. Human pseudotemporal expression patterns of neuronal calcium channels CACNA1C, and GRIA2. f. Relative proportions of subcategories of neural activity genes present in human gene clusters 7 and 3. g. UMAP embedding of the mouse brain organoid dataset subsetted with cortical neurogenesis clusters in (Fig 1e). h. Heatmap of representative marker genes of clusters in (g). i. Pseudotime trajectories identified in the mouse RNA-seq dataset. j. Gene clustering based on differential expression patterns along the mouse cortical neurogenesis pseudotime trajectory. k. Dotplot of GO term enrichment analysis of gene clusters in (i). l. Mouse pseudotemporal expression patterns of Sox2 (neural progenitors), Eomes (intermediate progenitors), Marcks (axon initial segment), Camk2b (synapses) and Grin2a (mature synapses). m. Mouse pseudotemporal expression patterns of neuronal calcium channels Cacna1c, and Gria2. n. Relative proportions of subcategories of neural activity genes present in human gene clusters 7 and 3. o. Average expression profile of axonogenesis and neuronal calcium channel genes, with shade representing standard deviations, along aligned pseudotime trajectories in human and mouse. ChP, choroid plexus. Rec., receptors. Glut., glutamate. Syn., synaptic. NT, neurotransmitter. RG, radial glia. IPs, intermediate progenitors. CNs, cortical neurons. Norm., normalized.

Unsupervised clustering of differentially expressed genes along the cortical neurogenesis trajectory revealed seven clusters with distinct expression dynamics (Fig 2b). Clusters 0, 1, 2, 4, 5, and 6 comprised genes displaying a decreasing pseudotemporal expression. These clusters were enriched for GO terms related to the cell cycle as well as more general cellular functions, and included neural progenitor genes such as SOX2, PAX6, and EOMES, reflecting the transition from neurogenesis to postmitotic neuronal development (Fig 2c,d, Extended Data Fig 3a, Supplementary Table 1). Clusters 3 and 7 represented genes with increasing pseudotemporal expression following distinct patterns (Fig 2b). The expression profile of genes in cluster 3 were distinguished by a progressive increase with a relatively late expression onset (Fig 2b). These genes were enriched for GO terms related to synapse formation and function and included synaptic genes such as CAMK2B, SHANK3, and the relatively late maturation synaptic gene GRIN2A (Fig. 2c,d, Extended Data Fig 3a, Supplementary Table 1). In contrast, genes in cluster 7 exhibited an earlier expression onset with an accelerated increase at early stages followed by a more subtle increase at later stages. As expected for this expression pattern, this cluster was enriched for GO terms associated with axon development and included axonal genes such as GAP43, NFASC and CNTN2 (Fig 2b-d, Extended Data Fig 3a, Supplementary Table 1). At the protein level, similar temporal dynamics were found for markers of neuronal development (Extended Data Fig 3b,c). Thus, cluster 7 represents the axonogenesis program, while cluster 3 represents the start of synaptogenesis.

We hypothesized that regulators controlling the timing of neurodevelopmental transitions, such as those from axonogenesis to synaptogenesis, should be present prior to the transition and would therefore most likely reside within cluster 7. A notable observation when analysing genes in cluster 7 was that these, in addition to axon development, were also strongly enriched for GO terms related to synapses (Fig 2c), and among them were synapse and neural activity genes such as GRIA2, GRIA3 and CACNA1C (Fig 2e). Indeed, the expression profile of these genes appeared more similar to axon outgrowth and axon initial segment genes than other synapse genes, suggesting that they were expressed before the start of synaptogenesis, and maintained consistently high expression levels during subsequent development (Fig 2d,e, Extended Data Fig 3d). This was confirmed by analysis of *in vivo* RNA sequencing data from the human prefrontal cortex, which showed not only similar developmental expression patterns of these genes but also confirmed that the onset of their expression precedes the onset of the synaptogenesis window, which starts around 27 weeks postconception (Extended Data Fig 3e) ^29^. Further analysis of this group of neural activity and synapse genes in cluster 7 revealed that their identities are biased toward neuronal calcium channels, as well as synaptic scaffold proteins which are known to be important for anchoring ion channels at their respective locations at the plasma membrane (Fig 2f).

Performing the same pseudotime analysis on the mouse scRNA-seq dataset, subsetted for neuronal progenitor and neuron clusters (Fig 2g,h), revealed a highly similar pattern. We identified a pseudotime trajectory reflecting cortical neurogenesis, as well as a trajectory corresponding to the early stages of gliogenesis (Fig 2i). Analysis of gene expression dynamics along the cortical neurogenesis trajectory revealed six clusters with decreasing pseudotemporal expression (1, 2, 3, 4, 5, 7, 8) which, based on GO term enrichment analysis and presence of neural progenitor markers, reflected the transition from neurogenesis to postmitotic neuron development (Fig 2j-l, Extended Data Fig 3f, Supplementary Table 2). Clusters 6 and 0 displayed increasing expression along pseudotime and reflected axon development and synaptogenesis, respectively, with their gene identities, expression profiles, and GO term enrichments similar to those of the corresponding human clusters (Fig 2j-l, Extended Data Fig 3f). Moreover, similar to the human data, cluster 6 was also associated with synapse-related enriched GO terms and included neural activity genes displaying early expression dynamics, with a bias toward neuronal calcium channels (Fig 2k-n, Extended Data Fig 3f,g).

A notable difference when comparing the pseudotemporal dynamics of the early expressed neuronal calcium channels between human and mouse, was their expression profiles relative to those of other axon development genes (Fig 2o). In the mouse dataset, the identified neuronal calcium channels had a pseudotemporal expression pattern preceding axon development genes, whereas expression profiles of these same genes were delayed in human along aligned pseudotime trajectories (Fig 2o, Extended Data Fig 4a,b). The species-specific differences in calcium channel expression profiles beyond the general delay in other developmental genes, combined with the early onset and sustained expression profile of these calcium channels, as well as the well-established roles of calcium dynamics in neurogenesis and neuronal morphogenesis ^40,64–73^, led us to investigate whether calcium dynamics might play a role in setting the species-specific cell state durations during early neuronal development.

### Species-specific differences in calcium dynamics at stage-matched timepoints

The delayed shift of neuronal calcium channel expression profiles relative to other developmental marker genes in human compared to mouse suggests that calcium dynamics may be regulated differently at matched stages across species. We used the GCaMP7f reporter to examine axonal calcium dynamics across developmental stages as neural activity modes change. Immature neurons initially display spontaneous calcium transients that gradually transition to synchronous network bursts, before the shift to asynchronous networks that start as synaptic structures begin to increase (Extended Data Fig 4c) ^74–76^. In both human and mouse organoids, we observed spontaneous calcium transients during early neuronal development, with the appearance of synchronous calcium network bursts increasing over time (Fig 3a-f, Extended Data videos 1-4). The transition from spontaneous to synchronous activity occurred around day 76 in human organoids and day 17 in mouse organoids (Fig 3e,f), which also corresponded with the onset of gliogenesis in each species (Extended Data Fig 4d-f), providing a carefully stage-matched timepoint for subsequent analyses (Fig 3g). To identify the timepoints reflecting the start of asynchronous networks, we assessed the number of synapses over time, as measured by co-localisation of pre- and postsynaptic puncta (Extended Data Fig 4g-i), which gives a more reliable read-out compared to analysing individual puncta (Extended Data Fig 4j-m). Similar to *in vivo*, a few synapses were already present during early stages, but only began to significantly increase at day 146 in human ALI-COs and day 28 in mouse ALI-COs (Extended Data Fig 4h,i), indeed substantially later than the onset of synchronous network burst activities (Fig 3e,f).

**Figure 3:**
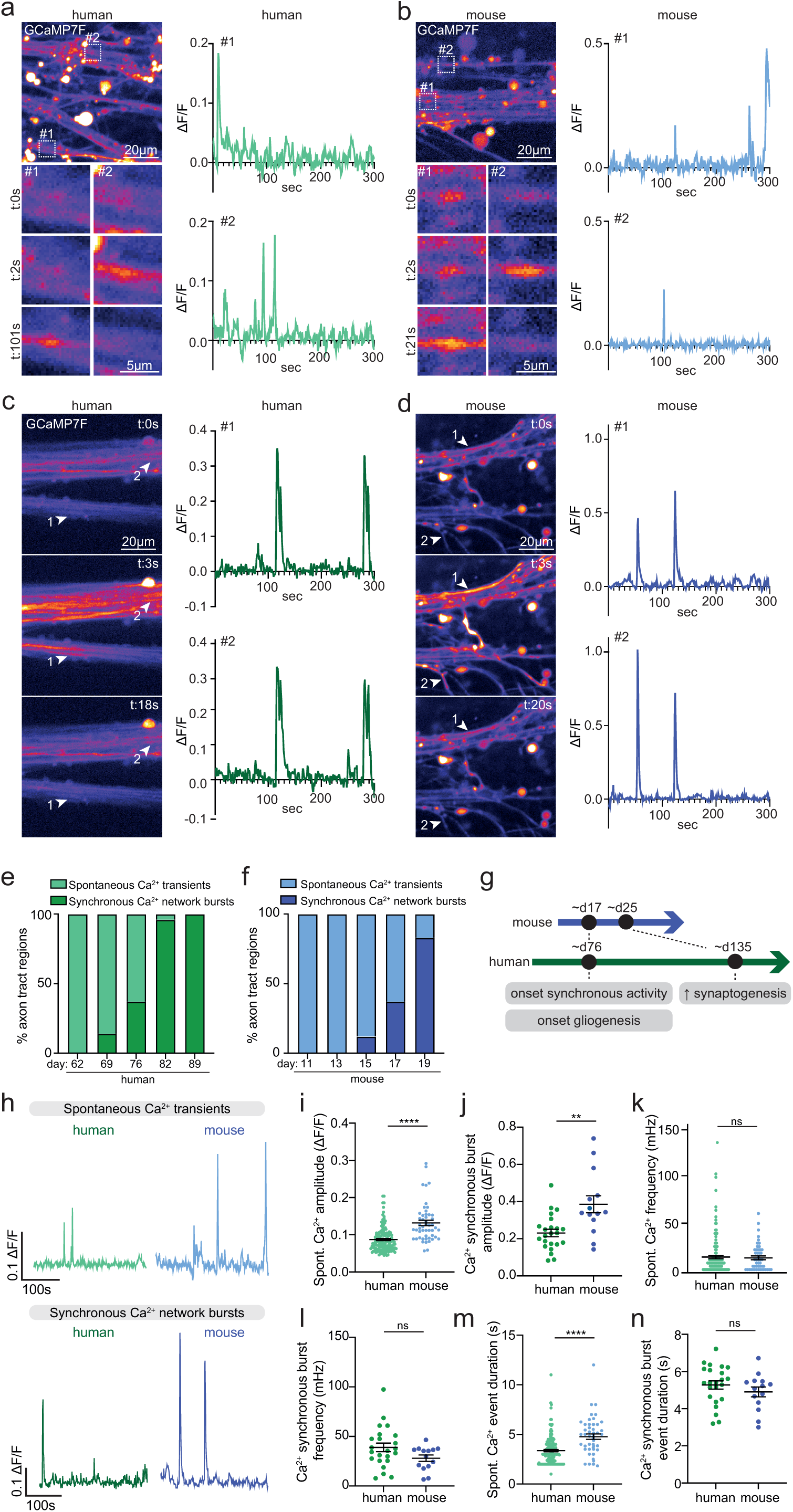
Species-specific differences in cytoplasmic calcium influx at stage-matched timepoints during spontaneous and synchronous activity. a. Axon tract of a human (day 69) brain organoid slice in micropatterned devices expressing GCaMP7f. Zooms show representative timelapse of spontaneous calcium transients at different timepoints in individual axons, with ΔF/F0 fluorescent GCaMP7f intensity plots of axons in zooms. b. Axon tract of a mouse (day 15) brain organoid slice in micropatterned devices expressing GCaMP7f. Zooms show representative timelapse of spontaneous calcium transients at different timepoints in individual axons, with ΔF/F0 fluorescent GCaMP7f intensity plots of axons in zooms. c. Timelapses of an axon tract of a human (day 81) brain organoid slice in micropatterned devices expressing GCaMP7f, showing synchronous calcium activity, with ΔF/F0 fluorescent GCaMP7f intensity plots of axons indicated by arrowheads. d. Timelapses of an axon tract of a mouse (day 19) brain organoid slice in micropatterned devices expressing GCaMP7f, showing synchronous calcium activity, with ΔF/F0 fluorescent GCaMP7f intensity plots of axons indicated by arrowheads. e. Quantifications of the relative proportion of axon tract regions of human brain organoid slices in micropatterned devices exhibiting spontaneous or synchronised calcium activity at different timepoints. Day 62: n=6 regions of interest, Day 69: n=34, Day 76: n=29, Day 83: n=12. f. Quantifications of the relative proportion of axon tract regions of mouse brain organoid slices in micropatterned devices exhibiting spontaneous or synchronised calcium activity at different timepoints. Day 11: n=7 regions of interest, Day 13: n=21, Day 15: n=16, Day 18: n=6, Day 20: n=10, Day 24: n=3. g. Schematic of timepoints of developmental milestones in human and mouse brain organoids, indicating day 17 in mouse and day 76 in human as stage-matched timepoints based on data shown in Figure 3f and Extended Data Figure 4b-k. h. GCaMP7f intensity plots of axons of a human (day 69) and a mouse (day 15) brain organoid slice culture in micropatterned devices at stage-matched timepoints during spontaneous calcium activity (day 69 human, day 15 mouse) and synchronous calcium network activity (day 82 human, day 19 mouse). i. Quantifications of average mean amplitudes of spontaneous calcium transients in human (day 69) and mouse (day 15) brain organoid slice cultures in micropatterned devices expressing GCaMP7f. Human: n=151 axons, mouse n=50. j. Quantifications of average mean amplitudes of synchronous calcium bursts in human (day 82) and mouse (day 19) brain organoid slice cultures in micropatterned devices expressing GCaMP7f. Human: n=23 regions of interest (average of 3 axons per ROI), mouse n=14. k. Quantifications of average frequencies of spontaneous calcium transients in human (day 69) and mouse (day 15) brain organoid slice cultures in micropatterned devices expressing GCaMP7f. Human: n=151 axons, mouse n=50. l. Quantifications of average frequencies of synchronous calcium bursts in human (day 82) and mouse (day 19) brain organoid slice cultures in micropatterned devices expressing GCaMP7f. Human: n=23 regions of interest (average of 3 axons per ROI), mouse n=14. m. Quantifications of average event durations of spontaneous calcium transients in human (day 69) and mouse (day 15) brain organoid slice cultures in micropatterned devices expressing GCaMP7f. Human: n=151 axons, mouse n=50. n. Quantifications of average event durations of synchronous calcium bursts in human (day 82) and mouse (day 19) brain organoid slice cultures in micropatterned devices expressing GCaMP7f. Human: n=23 regions of interest (average of 3 axons per ROI), mouse n=14. Spont., spontaneous. All data are shown as mean ± SEM; *ns P* > 0.05, \*\**P* < 0.01, \*\*\*\**P* < 0.0001 as determined by unpaired *t* tests.

We then performed a cross-species analysis of calcium dynamics at matched timepoints during spontaneous and synchronous activity. During both developmental neuronal activity modes, we observed a consistent decrease in amplitude, but not frequency, in human compared to mouse axons (Fig 3h-l). Calcium transient durations were also shorter in human during spontaneous activity, with no differences during synchronous burst activity (Fig 3m,n). We also compared proximal, mid and distal regions of axon tracts and observed very similar parameters, suggesting a consistent species-specific difference irrespective of location on the axon tract (Extended Data Fig 4n-s). The identified species-specific differences in calcium dynamics displayed a quantitative difference even at matched developmental stages. This suggests that calcium dynamics not only correlate with species-specific developmental timing but represent a strong candidate mechanism for setting developmental timing.

### L-type VGCC-mediated calcium influxes modulate developmental transition timing and axon tract length

Next, we sought to investigate if species-specific calcium dynamics could be responsible for setting developmental timing in a manner leading to their respective differences in axon tract length. To address this, we aimed to identify the calcium channels driving calcium dynamics during these neuronal development stages. The scRNA-seq analysis revealed an early onset of expression of plasma membrane-localized calcium channels in both human and mouse neurons. Additionally, the observed calcium dynamics were characterized by spikes, which typically result from extracellular calcium influx through plasma membrane-localized channels rather than release from intracellular calcium stores that typically lead to calcium waves, suggesting these calcium channels could be responsible ^65,66^. Moreover, the long duration of the observed calcium transients, lasting several seconds, pointed to calcium channels with a slow inactivation rate, which is typical for the L-Type VGCCs, one of the channel types we found to have an early onset expression profile in scRNA-seq ^77^. We therefore tested the effects of treatment with an L-Type VGCC antagonist, which we found to inhibit nearly all calcium dynamics in early stages corresponding with spontaneous activity, as well as during later stages corresponding to synchronous activity, and could be recovered upon washout (Fig 4a,b, Extended Data Fig 5a, Supplementary videos 5-10). Instead, treatment with an L-type VGCC agonist led to a profound increase in calcium amplitudes, similar to those observed in untreated mouse organoids, with a subtle decrease in frequency and no difference in event duration or number of axons firing (Fig 4a-c, Extended Data Fig 5b,c).

**Figure 4:**
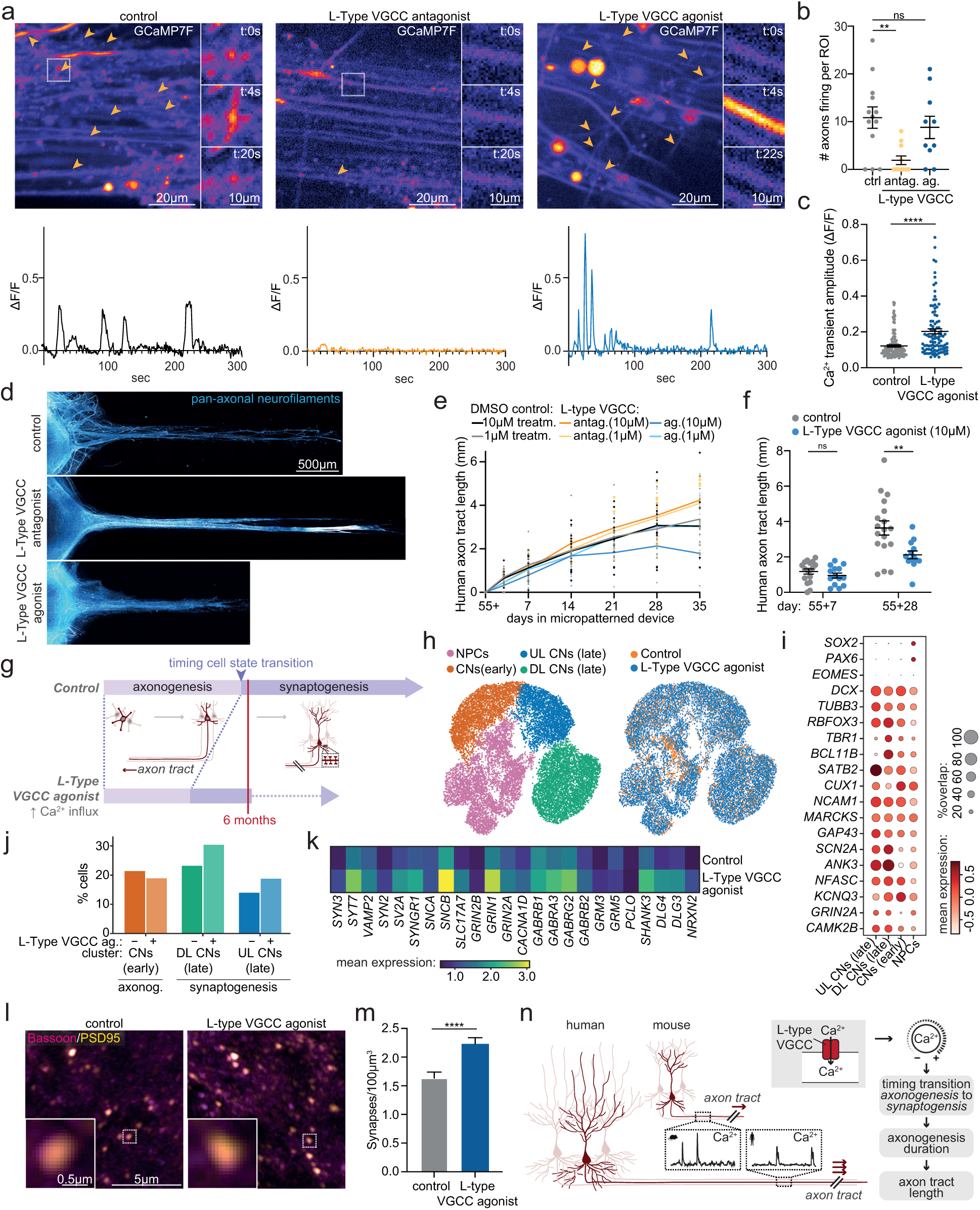
L-type VGCC-mediated calcium dynamics set the timing of the transition from axonogenesis to synaptogenesis. a. Axon tract of human (day 70) brain organoid slices in micropatterned devices expressing GCaMP7f and treated with DMSO (control), nimodipine (L-Type VGCC antagonist) or BayK8644 (L-Type VGCC agonist). Arrowheads mark axons showing calcium transients during a five minute timelapse, indicating nearly complete perturbation of calcium dynamics upon L-Type VGCC blockage. Zooms show representative timelapse of individual axons, with ΔF/F0 fluorescent GCaMP7f intensity plots of axons in zooms. b. Quantifications of number of axons firing in an ROI during a five minutes timelapse in axon tracts of human brain organoid slices (day 70) in micropatterned devices expressing GCaMP7f and treated with DMSO (control), nimodipine (L-Type VGCC antagonist) or BayK8644 (L-Type VGCC agonist). Control: n=13 regions of interest, L-Type VGCC antag.: n=11, L-Type VGCC ag.: n=10. c. Quantifications of average mean amplitudes of spontaneous calcium transients in human brain organoid slice cultures (day 70) in micropatterned devices expressing GCaMP7f and treated with DMSO (control) or BayK8644 (L-Type VGCC agonist). Control: n=137 axons, L-Type VGCC ag.: n=100. d. Representative images of axon tracts of human (day 90) brain organoid slices in micropatterned devices immunostained for pan-axonal neurofilaments and treated with DMSO (control), nimodipine (L-Type VGCC antagonist) and BayK8644 (L-Type VGCC agonist). e. Quantifications of axon tract length of human brain organoid slice cultures in micropatterned devices, treated with DMSO (control), nimodipine (L-Type VGCC antagonist) or BayK8644 (L-Type VGCC agonist), at different timepoints. n=8-32 organoids. Control : n=19 organoids (1 µM) and n=12 (10 µM), L-Type VGCC antag.: n=12 (1 µM) and n=9 (10 µM), L-Type VGCC ag.: n=15 (1 µM) and n=9 (10 µM). Mixed-effects model with Geisser-Greenhouse correction P<0.0001 for time, p=0.0219 for treatments. f. Quantifications of axon tract length of human brain organoid slice cultures in micropatterned devices, treated with DMSO (control) or BayK8644 (L-Type VGCC agonist), at day 55+7 and 55+28. Control: n=15 organoids (55+7) and n=14 (55+28), L-Type VGCC ag.: n=11 (55+7) and n=8 (55+28). Multiple unpaired t-test with two-stage step-up (Denjamini, Krieger and Yekutieli; FDR 5%). g. Schematic model of an earlier cell state transition from axonogenesis to synaptogenesis upon L-type VGCC agonist treatment, suggesting that neurons spend a relatively longer time in the synaptogenesis window after 6 months in culture. h. UMAP embedding of day 90 human mouse brain organoids treated with DMSO (control) or BayK8644 (L-Type VGCC agonist), coloured by Leiden clusters or treatment. i. Dotplot of representative marker genes of clusters in (h). j. Relate cell proportion normalized per condition across the cortical neuron clusters in (l) undergoing axonogenesis (early) or synaptogenesis (late). k. Heatmap of synapse gene expression of non-zero expressing cells in upper layer neuron cluster in (h), comparing control and L-type VGCC agonist conditions. l. Representative images of 6 months old human brain organoid slice cultures, treated with DMSO (control) or BayK8644 (L-Type VGCC agonist), and immunostained for bassoon and PSD95. Zoom represents co-localisation of pre- and postsynaptic structures. m. Quantifications of number of co-localising bassoon and PSD95 puncta in 6 months old human brain organoid slice cultures, treated with DMSO (control) or BayK8644 (L-Type VGCC agonist). DMSO: n=150 regions of interest, L-Type VGCC ag.: n=182. n. Schematic of model. ROI, region of interest. Antag., antagonist. Ag., agonist. Treatm., treatment. CNs, cortical neurons. DL, deep layer. UL, upper layer. Axonog., axonogenesis. All data are shown as mean ± SEM; ns *P/Q* > 0.05, \**P* < 0.05, \*\**P* < 0.01, \*\*\*\**P* < 0.0001 as determined by unpaired *t* tests unless stated differently.

We then examined the effect of these perturbations on human axon tract length, to test if this could be influenced by increasing or decreasing calcium dynamics. We observed that while potentiation of calcium amplitude, thereby mimicking mouse calcium dynamics, did not affect initial axon outgrowth rates, with similar axon lengths observed after 7 days, it did result in a shorter axonogenesis time window which, similar to mouse, led to shorter axon tracts (Fig 4d-f). On the other hand, blocking L-type VGCC-mediated calcium influxes resulted in a trend towards a longer axon outgrowth time window and an increase in axon tract length, beyond the unperturbed human axons (Fig 4d-e).

In mature neurons, the relatively long-lasting calcium influxes mediated by L-type VGCCs typically lead to more long-term modulatory responses, aligning with our findings of its effect on axon tract development duration rather than outgrowth rates. An important pathway for these modulatory responses involves downstream increases of the second messenger cAMP upon L-Type VGCC-mediated calcium influxes, with cAMP in turn also regulating L-Type VGCC activity, which has been linked to neuronal development and plasticity and typically involves changes in gene expression ^78–81^. We therefore tested whether cAMP similarly impacted axon tract growth length and timing. Brain organoids treated with cAMP during axon outgrowth displayed truncated axon tract growth duration and shorter axon tracts, with no changes in axon tract growth rates whilst in their elongation stage, thus phenocopying the effects observed with an increased L-Type VGCC-mediated calcium influx (Extended Data Fig 5d-f).

These findings indicate that species-specific calcium differences, through L-Type VGCCs and cAMP, regulate axon tract growth duration, potentially through modulatory responses involving regulation of gene expression. To examine the downstream effects on gene expression, we performed single cell RNA sequencing analysis of human organoids treated with DMSO or an L-Type VGCC agonist, the condition that mimicked mouse calcium dynamics (Fig 4c). We analysed organoids at 6 months to capture the effect on transition timing from axonogenesis to synaptogenesis (Fig 4g) since at this timepoint a subset of the human neurons have transitioned to synaptogenesis, starting around 5 months (Extended Data Fig 4h). Cluster analysis revealed the expected stages of cortical neurogenesis, including a cluster representing neural progenitor cells, and several clusters representing neurons at different stages (Fig 4h,i). These included deep and upper layer neurons still undergoing axonogenesis, deep layer neurons undergoing synaptogenesis and upper layer neurons undergoing synaptogenesis. Of note, in L-Type VGCC agonist-treated organoids there were relatively more cells in synaptogenesis clusters and fewer in the axonogenesis cluster, indeed pointing to an earlier transition (Fig 4j).

We then examined the expression levels of synapse genes, focusing specifically on those in the previously identified gene cluster 3, which, unlike the synaptic genes in cluster 7, likely reflect actual synaptogenesis (Fig 2b,c, Supplementary Table 1). In both the more mature deep and upper layer neuron clusters, we observed a global increase in synapse gene expression in non-zero-expressing cells upon L-type VGCC agonist treatment (Fig 4k; Extended Data Fig 5g,h). Consistent with this, we also detected higher synapse numbers at similar timepoints following L-type VGCC agonist treatment (Fig 4l,m), confirming the earlier transition to synaptogenesis. Together, these data show that human axons, compared with those of mice, exhibit reduced calcium influxes, which extends axon growth duration by delaying the transition to synaptogenesis, thereby giving rise to longer axon tracts in humans (Fig 4n).

## DISCUSSION

We demonstrate a causal relationship between species-specific developmental timing and morphology, specifically axon length, driven by a selective modulation of developmental duration. By focusing on axon tract development, as opposed to later neuronal morphogenic stages of dendritogenesis and synaptogenesis, we could assess the complete developmental process within a realistic timeframe of months instead of years. We generated modified human and mouse brain organoid slice cultures that extended self-organised axon tracts into channels of micropatterned devices. Investigating axon development in the tissue context is relevant, given the differences in the developmental onset and properties of synchronous networks as well as axon growth cone behaviours seen in 2D cultures ^82,83^. While the exact topology of organoids still diverges from the *in vivo* brain, the methodologies described here enabled axon tract development within a more physiological environment than 2D cultures and, combined with the micropatterned devices, showed similarity to commonly used *ex vivo* cultures for axon growth studies ^84–86^.

Using these integrated methods, we identified a cell-intrinsic species-specific difference in the duration of axon growth that leads to increased length of human axons. The finding that growth duration is independent of tissue environment, together with extensive previous work on axon pathfinding, suggest multiple layers of control in establishing axon tract length: a species-specific time window, combined with well-described interactions with environmental factors like guidance molecules and stiffness ^55,56^. We also observed a slower rate of growth in humans, in line with previous studies that have identified roughly two-fold slower rates of biogenesis and metabolism ^35–38^. However, while previously identified changes in tempo explain allochronic changes in timing ^87^, they do not explain differences in size, since slower growth rate even if coupled with a proportionate increase in duration would not lead to greater growth. Instead, we uncovered a mechanism involving tuning of calcium and cAMP levels to set developmental duration independently of rate, by controlling the timing of the transition from axonogenesis to synaptogenesis.

Whilst calcium dynamics had not yet been linked to evolutionary timing of brain development, perturbations in spontaneous activity have been found to lead to mistargeted or aberrant axon projections, consistent with our results ^88,89^. Spontaneous calcium transients also mediate remodelling of the axonal cytoskeleton, thereby controlling transient processes such as axon turning and growth cone pausing ^67,70,71,73^. However, the mechanism we identified in calcium-mediated development of species-specific axon tract lengths seems not to be driven by transient processes, as we found no differences in axon tract growth rate upon perturbation of L-type VGCC-mediated calcium dynamics. These perturbations were accompanied by changes in synaptic gene expression, consistent with the well-described role of calcium and cAMP dynamics on gene expression in mature neurons, particularly in the context of synaptic plasticity ^90–93^. It would be interesting to direct future research towards investigating how the cumulative effect of species differences in calcium influx may influence genetic developmental programs.

Evolutionary mechanisms ultimately stem from genetic differences, and it is unknown what genetic changes might be responsible for the species-specific differences in calcium dynamics and expression profiles of neuronal calcium genes. Relevant genetic changes could affect calcium channel expression levels, their targeting to plasma membranes or their turnover. These genetic changes may not be limited to changes in L-Type VGCC channels, as we observed a common signature of calcium-permeable ion channels with an early expression onset. An interesting avenue would be to explore the effect on developmental duration of other channels, such as NMDA receptors that were recently identified to exhibit calcium-mediated roles in synapse and dendrite formation ^64^.

The scRNA-seq data also pointed to a pattern of calcium channels expressed even earlier, during neurogenesis and neuron development, suggesting a potential role of calcium-mediated regulation of developmental transition timings of other neurodevelopmental processes. Consistent with this, calcium activity is involved in many processes of neuron development and plasticity, with perturbations of calcium dynamics affecting neurogenesis, axon pathfinding as well as regeneration, synapse formation and synaptic plasticity ^40,64–73^. Impairments in calcium dynamics have also been linked to various neurodevelopmental disorders, including epilepsy, intellectual disability and autism spectrum disorder ^94–97^.

Timothy syndrome is specifically associated with predicted gain-of-function mutations in L-Type VGCCs, and its dysregulation is linked to cognitive impairments and intriguingly, microcephaly, or smaller brain size, which aligns with our observations of decreased growth ^98,99^. Thus, the widespread role of calcium activity throughout neurodevelopment hints at a broader regulatory function of calcium dynamics in coordinating the timing of neurodevelopmental transitions, pointing to an evolutionary timing mechanism that may allow neurons with distinct species-specific complexities beyond axon tracts to arise. Finally, as is common in evolution, a calcium and cAMP-mediated mechanism of timing is likely not a new innovation of the human lineage, and indeed, oscillatory cAMP signalling acts as a developmental timer in the amoeba species *D. discoideum* ^100^. Given this early evolutionary emergence, as well as the appearance of calcium dynamics and ubiquitous roles of cAMP across cell types and species, it would be interesting to explore if this timing mechanism also drives other developmental processes, even beyond neurodevelopment.

## MATERIAL & METHODS

### Cell lines

The human H9 (WA09, female) and H1 (WA01, male) embryonic stem cell lines were purchased from WiCell. The U.K. Stem Cell Bank Steering and the ERC ethics committee has approved the use of human embryonic stem cells for this project, and the cell lines are registered on the Human Pluripotent StemCell Registry (hpscreg.eu). The mouse E14 embryonic stem cell line (ESCs; RRID: CVCL_C320) was kindly gifted by the group of Marta Shahbazi. Human embryonic stem cells were maintained in StemFlex (Thermo Fisher Scientific, A3349401). Unless specified otherwise, human organoids were generated from H9 stem cells. E14 embryonic mouse stem cells were expanded in 2iLIF media and transferred to StemFlex supplemented with 1000 U/ml ESGRO recombinant mouse leukemia inhibitory factor (mLIF; Sigma-Aldrich #ESG1107) before generation of organoids using the same protocol as for human, as described before ^47^. All stem cell lines were kept as feeder-free cultures on plates coated with growth factor reduced Matrigel (Corning #356230) and passaged about 2x a week using EDTA.

To generate the mCherry-expressing H1 and E14 lines, annotated as H1 PB001 and E14 PB001, plasmids PB001 carrying the mCherry cassette and pBase carrying the transposase (2 μg of each plasmid) were electroporated into 10 million H1 or E14 cells using the Human Stem Cell Nucleofector Kit 1 (Lonza, VPH-5012). Following electroporation, cells were grown in one well of a 6-well plate in StemFlex supplemented with 1 nM ROCK inhibitor (BD Biosciences, 562822). After 48 hours post electroporation, the cells were selected using StemFlex supplemented with Puromyocin (Gibco, A11138-03) at a final concentration of 0.5 µg/mL.

### Plasmids

The pGP-AAV-syn-jGCaMP7f-WPRE plasmid was obtained from Douglas Kim and the GENIE project (Addgene, 104488) ^101^. The MARCKS-GFP plasmid was obtained from the group of Casper Hoogenraad, with the cassette composed of the first 41 amino acids of the MARCKS protein containing an Alanine to Cysteine mutation (MGCQFSKTAAKGEAAAERPGEAAVASSPSKANGQENGHVKV), fused to a GS linker and TagRFP-T ^102^. The PB001 insertion plasmid contained the insert encoding a human ubiquitin promoter driving the expression of mCherry followed by a P2A ribosome skipping sequence and a puromycin resistance cassette, with a bGH polyA termination signal, flanked by PB transposase inverted terminal repeat sequences.

### Human and mouse brain organoids

Human and mouse cerebral organoids were generated, with some modifications, as described before, with both protocols following the same steps but with the timing of each step tailored to match their species-specific developmental tempo (Fig 1b) ^47,52,103^. Briefly, on day 0, human or mouse embryonic stem cells were dissociated with Accutase (Gibco, A1110501), resuspended in EB Formation medium (StemCell Technologies, 8570) supplemented with 50 μM ROCK inhibitor Y27632 (SantaCruz Biotechnology, sc-281642A) and 4000 cells per well were seeded into ultra-low attachment U-bottom 96-well plates (Corning, 7007) to form embryoid bodies. On day 5 for human or day 2 for mouse, the media was replaced with Neural Induction medium (StemCell Technologies, 8570). On day 7 for human or day 5 for mouse, embryoid bodies were embedded in Matrigel (Corning, 354234) and transferred to 6-well plates with Expansion medium (StemCell Technologies, 8570). On day 10 for human and day 6 for mouse, the culture media was replaced with Improved Differentiation Medium without vitamin A (IDM-A: 1:1 [v/v] DMEM/F12 (Gibco, 11330032) and Neurobasal (Gibco, 21103049) media, 1:200 [v/v] N-2 supplement (Gibco, 7502048), 1:50 [v/v] B-27 supplement minus vitamin A (Gibco, 12587010), 1x GlutaMAX (Gibco, 35050038), 0.5x MEM non-essential amino acid solution (MEM-NEAA; Gibco, M7145), 2.5 μg/mL human insulin (Sigma-Aldrich, I9278), 50 μM 2-mercaptoethanol (Gibco, 31350010), 1:100 [v/v] penicillin-streptomycin (Gibco, 15140122)). On day 14 for human or day 7 for mouse, Matrigel was removed from the organoids, by either manual dissection or chemical Matrigel removal. For the latter, the bulk of Matrigel was initially removed by manual dissection, after which organoids were transferred to a dish, covered with cell recovery solution (Corning #354253) and incubated at 4°C for 20 min with regular gentle agitation. Cell recovery solution was removed and organoids were gently washed thrice with phosphate-buffered saline (PBS; 125 mM NaCl, 17 mM Na2HPO4, 9 mM NaH2PO4, pH 7.4). Organoids were kept on an orbital shaker (57 RPM, 25 cm orbit) and cultured in 6 cm dishes in IDM-A medium until day 30 for human or day 8 for mouse, and then in IDM-A medium supplemented with 1:50 [v/v] Matrigel until ready for slicing.

### Human and mouse brain organoid slice cultures

Human and mouse brain organoids were sliced similar as previously described, with some modifications ^48^. In brief, unless stated differently, mouse and human organoids were harvested on respectively day 8 and 55. Organoids were embedded in 3% low-gelling temperature agarose (Sigma, A9414) in molds (Sigma, E6032), incubated on ice for 30-60 min, and sectioned at 300 µm thickness using a Leica VT1000S vibrating microtome. Organoids slices were kept as ALI-COs or transferred to micropatterned devices for axon outgrowth experiments. For ALI-CO cultures, slices were cultured on Millicell-CM culture inserts (Millipore, #PICM0RG50). For axon tract outgrowth assays, slices were transferred to micropatterned devices as detailed below. Brain organoid slice cultures were kept in serum-free slice culture medium (SFSCM; Neurobasal (Invitrogen, #21103049), 1:50 (v/v) B27+A (Invitrogen, cat. #17504044), 1X (v/v) Glutamax (Gibco #35050038), 1:100 [v/v] penicillin-streptomycin (Gibco), 1:500 (v/v) Fungin (Invivogen, #ant-fn)). After 14 days for human or 3 days for mouse, and every 3-4 days after, ∼50% of the medium was replaced with BrainPhys medium (BP; BrainPhys Neuronal Medium N2-A SM1 kit (StemcellTechnologies, #05793), 1:100 [v/v] penicillin-streptomycin (Gibco), 1:500 (v/v) Fungin (Invivogen, #ant-fn)).

### Fabrication of micropatterned devices

Microfluidic devices were produced by replica molding using polydimethylsiloxane (PDMS, Sylgard 184; Ellsworth Adhesives, Wilmington, MA, USA) on a master mold that was fabricated on a silicon wafer using soft lithography ^104,105,106,107^. The master mold was fabricated to 450 μm thickness using SU8-100 photoresist according to the manufacturer’s (Kayaku® Advanced Materials) instructions and the OAI Model 204 Mask Aligner for UV exposure (Extended Data Fig 2b). The dimensions and surface characterization of the master mold are shown in Extended Data Fig 2d. Next, the PDMS pre-polymer and curing agent were mixed in a 10:1 weight ratio to make PDMS and then degassed in a desiccator before pouring onto the master mold. The PDMS was cured at 65°C for 8 hr (Extended Data Fig 2c). After curing, milli-wells were punched using a 3 mm biopsy puncher. Micropatterned devices were then soaked in ethanol, placed onto the glass bottoms of either 35 mm dishes (World Precision Instruments, D35-100) for live-imaging experiments, or 24-well plates (Cellvis, P24-0-N) or 12-well plates (Cellvis, P12-1.5H-N) for axon outgrowth assays, and kept in the hood overnight, allowing evaporation of ethanol and thereby enabling bonding.

### Brain organoid slice cultures in micropatterned devices

The chambers and channels of the micropatterned devices were coated with 0.5 mg/mL growth factor reduced Matrigel (Corning, 356230) in DMEM/F12 for at least 1h at 37°C/5%CO2. Just before placing brain organoid slices (generated as described above) in the micropatterned devices, Matrigel coating was removed, and brain organoid slices were gently placed in the chambers directly adjacent to the opening of the channel. Any reminiscent liquid was removed and slices were kept at 37°C/5%CO2 for 30-60 min after which SFSCM medium was added. Subsequent medium changes were the same as with the ALI-CO cultures.

### Mouse spinal cord explant co-culture in micropatterned devices

Mouse tissue was collected in accordance with the UK Animals (Scientific Procedures) Act 1986 and European Community Council Directive on Animal Care by a trained technician of the MRC-LMB animal facilities. Mouse spinal cord explants from C57BL/6 E12.5 embryos were obtained as described before ^48^. In brief, mouse uteruses and subsequently dissected embryos were handled in ice-cold HBSS without Calcium and Magnesium (Gibco, 14175095) and kept on ice throughout the handling process. Using microdissection equipment, embryos were excised and liberated from the uterine horns, after being pinned down supine through their limbs for stability. The dorsal skin, meninges, and vasculature were peeled away from the back using fine tweezers to expose the spinal cord in the midline. Cuts were made 1 mm laterally down each side of the spinal cord, to free it from the sides of the body. Subsequently, the excised spinal cord and surrounding tissue was turned on its side so that the viscera could be removed from its ventral aspect, leaving just the spinal cord and some associated paraspinal muscle around it. The tail and head were removed from each respective end. The dissected mouse spinal cords were then sectioned, using the same protocol as described above, and slice cultures were placed in the channel of the micropatterned devices opposite the organoid slice.

### Pharmacological treatments

For drug treatments of brain organoid slices in micropatterned devices, one day after the slice cultures were placed in the devices, and during subsequent medium changes, medium was replaced with SFSCM or BrainPhys (at similar timepoints as with ALI-CO cultures described above) supplemented with indicated final concentrations of either 1 μM or 10 μM BayK8644 (Stratech Scientific, S7924-SEL), 1 μM or 10 μM Nimodipine (Selleckchem, S1747) or 100 μM 8-Br-cAMP (BioLog, B007). For the wash-out experiment in Extended Data Fig 5a, submerged brain organoid slices were incubated with BrainPhys supplemented with 10 μM Nimodipine (Selleckchem, S1747) for 1h and kept on an orbital shaker (57 RPM, 25 cm orbit). After timelapse acquisitions, medium was replaced with BrainPhys and slices were placed back on the shaker for 3h for the wash out experiment, followed by another series of timelapse acquisitions.

### Viral labelling

For sparse labelling of cells enabling assessing cell morphologies, ALI-COs were induced with 0.05 - 0.5 μl Sendai virus carrying emGFP (ThermoFisher, A16519) diluted in 2 μl SFSCM for mouse ALI-COs or 10 μl SFSCM for human ALI-COs that was added dropwise onto the slice. For MARCKS-GFP labelling of human brain organoid slices in co-culture with mouse spinal cord explant, human brain organoid slices in micropatterned devices were induced with lentivirus carrying doxycycline-induced MARCKS-GFP, generated as described in ^102^, ∼1-3h after being placed in the devices. For viral induction, medium was removed, and virus diluted in 10 μl SFSCM was added to the chamber, after 30-60 min a total of 2 ml SFSCM was added back again. The next day, slices were washed before mouse spinal cord explants were placed on the opposing chambers of the devices to ensure specific labelling of only human organoids. For calcium imaging, brain organoids in micropatterned devices were induced with AAV1 virus carrying GCaMP7f (Addgene, 104488), using the same viral induction protocol as described above, within 3-36h after organoids were placed in the micropatterned devices. Calcium imaging was performed at least 7 days after viral induction to ensure sufficient GCaMP7f expression.

### Fixation, cryosectioning and immunostaining

At indicated timepoints, ALI-COs were fixed overnight at 4 °C with 4% [v/v] paraformaldehyde in phosphate buffer (145 mM Na2HPO4, 21 mM NaH2PO4, pH 7.4) and washed thrice with PBS. Whole ALI-COs were incubated with primary antibodies and secondary antibodies for 48 h at room temperature, with 3x 8 h washes after every antibody incubation. Whole ALI-COs were mounted using Prolong Diamond (Invitrogen, P36970) with coverslips on microscope glass slides. For cryosectioned samples, fixed ALI-COs were incubated in phosphate buffer with 30% [w/v] sucrose at 4°C overnight. Organoids were embedded in 7.5% [w/v] gelatin (Sigma-Aldrich, G1890), 30% sucrose in phosphate buffer, and the gelatin blocks were frozen by submersion in cold isopentane (-50 °C) for 90 s. Frozen blocks were stored at -80 °C until sectioned in 20 μm slices using a CM1950 cryostat (Leica Biosystems), after slices were collected on charged slides (SuperFrost Plus Adhesion; Fisher Scientific, J1800AMNZ). Cryosections were incubated with primary antibodies overnight at 4°C, and secondary antibodies for 1h at room temperature, with 3x 10 min PBS wash steps after each antibody incubation. All primary and secondary antibodies were diluted in permeabilisation buffer (0.25% Triton-X, 4% donkey serum in PBS). The following primary antibodies were used in this study: Bassoon (Enzo Life Sciences, SAP7F407), GAPDH (Abcam, ab8245), GFAP (Abcam, ab7260), Homer1 (Synaptic systems, 16003), PSD95 (Abcam, ab18258), Pan-axonal neurofilaments Clone SMI312 (Biolegend, 837904), Synaptophysin (Abcam, ab32127), Trim46 (Synaptic Systems, 337003), VGAT (Synaptic Systems, 131003).

### Immunoblotting

At the time of sample collection, ALI-COs were cleared of residual agarose, washed thrice in ice-cold PBS, snap-frozen, and stored at -80°C. Protein lysates were obtained by thawing samples on ice, followed by homogenization by resuspending 20x with a Dounce homogenizer in 1 ml modified-RIPA (mRIPA: 1% Triton-X, 0.1% SDS, 150 mM NaCl, 50 mM Tris pH 7.4, 2 mM EDTA, 12 mM sodium deoxycholate) supplemented with protease inhibitors (Thermo Fisher, 78430), and incubated for 40 min at 4°C on a rotor. Protein lysates were centrifuged (20,000 *g*, 20 min, 4°C) and protein concentrations were obtained using Quick Start Bradford Dye Reagent (Bio-Rad, 5000205). Normalized protein samples were resolved on 4-20% tris-glycine gels and transferred to Amersham Hybond P 0.45 PVDF blotting membranes (GE Healthcare, 10600023). Membranes were blocked for 1h at room temperature in PBS-T with 5% milk, incubated overnight at 4°C with primary antibodies diluted in PBS-T with 2% milk, then for 1h at room temperature with secondary antibodies diluted in PBS-T with 2% milk, with 3x 10 min wash steps in PBS-T after each antibody incubation. Resolved proteins on the blots were visualised using a Gel Doc XR+ system.

### Microscopy, live-imaging and image analysis

Brightfield images of axon tracts growing in micropatterned devices and fluorescent images of the MARCKS-GFP-positive and mCherry-positive axon tracts for the co-culture experiments were acquired using an Evos XL Core microscope (Thermo Fisher Scientific). Images for synaptogenesis and gliogenesis were acquired on an inverted Zeiss LSM 880 Airyscan equipped with a 63x/1.4 NA Oil lens, and GaAsp, spectral and Airyscan detectors. For the synaptogenesis analysis, the super resolution settings of the Airyscan detectors were used to acquire high resolution images. Confocal imaging of all other fixed samples was performed on an inverted Nikon CSU-W1 spinning disk microscope, equipped with 10x/0.45 NA Air, 25x/1.05 NA Silicone Oil and 60x/1.2 NA Water lenses, and a sCMOS camera (95% QE). Displayed microscopy images represent maximum intensity projections of a Z-stack of the region of interest. Live-imaging of calcium dynamics was conducted on an inverted Nikon X1 Spinning Disk, containing a 60x/1.2NA Water lens, sCMOS cameras (95% QE), and a heated and gas-controlled incubation chamber. ALI-COs were kept at 37 °C and 5% CO2, and timelapses were acquired at a 1Hz imaging speed of a single plane. Image processing and analysis was performed in Fiji.

### Quantifications neuronal polarity, 3D neurite lengths and soma areas

Neurons were categorised as polarised when one of the neurites, the future axon, was at least two-fold longer than the second longest neurite. To assess neurite lengths, a line was manually drawn to trace the neurite length, a Z-plot was generated and the length of the neurite was traced to obtain 3D length measurements. Soma areas were obtained by manually tracing soma outlines.

### Quantifications of axon tract lengths in micropatterned devices

Brightfield images of the overlapping axon tract regions, covering the tract from beginning to end, were stitched using the Fiji plugin Pairwise Stitching ( https://github.com/fiji/Stitching/blob/Stitching_-3.1.9/src/main/java/plugin/Stitching_Pairwise.java) ^108^. Axon tract lengths were manually traced, using the surface of the organoid where the tract extends from as start and the most distal part of the tract, or individual pioneering axons, as end. For Fig. 4e, experiments were pooled, and because not all experiments contained every condition, we applied an inclusion criterion to the control samples. Specifically, controls were required to deviate by no more than 30% from the average control endpoint value. This was done to ensure a representative and comparable dataset across all conditions and treatments.

### Quantifications of proportion of spontaneous and synchronous calcium activity and subsequent stage-matching

Timelapses with 133.62×133.62μm image size and 5 min durations were acquired from axon tract regions in micropatterned devices. If at least two axons showed a clear synchronous firing pattern during the acquisition, it was classified as synchronous network burst activity, otherwise as spontaneous activity. Synchronous firing events first emerged on day 69 in human organoids and on day 15 in mouse organoids, while most calcium activity at these stages still consisted of non-synchronous spontaneous events. These matched timepoints were then used for cross-species analysis of spontaneous calcium activity, for which the only timelapses that did not display synchronous events were included. Cross-species analysis of synchronous burst activity was conducted on day 82 in human and day 19 in mouse, marking the first timepoints at which the majority of calcium activity events exhibited synchrony in both species. At these stages, robust burst activity was observed throughout the entire tract, whereas earlier timepoints showed that synchronous activity occasionally involved the coordinated firing of individual axons within the tract.

### Calcium activity quantifications

Regions of interest were manually delineated for each image. For spontaneous calcium activity measurements, all axons exhibiting calcium activity at least once during the acquired 5 min timelapses were traced and analysed. For synchronous calcium burst activity, three axons were traced per timelapse, and their read-outs were averaged and treated as a single data point. The corresponding temporal signal for each ROI was subsequently analyzed using a custom-developed ImageJ macro (https://github.com/MRC-LMB-Light-Microscopy-Facility/imagej-macro-calcium-imaging; DOI: 10.5281/zenodo.14562107). Briefly, the temporal signal was derived by calculating the mean pixel intensity within each ROI at each time point. To mitigate noise, the raw signal underwent sequential temporal filtering, employing a median filter followed by a minimum filter. Putative events were identified by detecting local maxima in the ratio exceeding a pre-defined threshold. Onset and offset times for each event were defined by contiguous temporal segments exceeding this threshold and containing a single local maximum. The spatiotemporal characteristics of each identified event, specifically its location and maximum amplitude, were recorded and subsequently aggregated for each ROI in order to compute mean amplitude and frequency. The ΔF/F traces were obtained using a sliding window of 21 frames. Calcium signals above a 0.045 threshold were identified as transients, followed by the analyses of mean amplitudes, event frequencies and event durations.

### Glia cell proportion quantifications

Images with a 66.10×66.10×1.5 μm volume of brain organoid cryosections immunostained for DAPI and GFAP were acquired, with regions selected using the DAPI channel to ensure blinded analysis. Nuclei numbers from a maximum intensity projection were assessed using the Fiji plugins Watershed Separation (https://github.com/imagej/ImageJ/blob/v1.54f/ij/plugin/filter/EDM.java) and Particle Analyzer (https://github.com/imagej/ImageJ/blob/v1.54f/ij/plugin/filter/ParticleAnalyzer.java), and then classified as GFAP-positive or GFAP-negative to calculate the respective cell proportions.

### Synapse quantifications

Images with a 19.44×19.44×1.35 μm volume of brain organoid cryosections immunostained for DAPI, Bassoon and PSD95 were acquired, with regions selected using the DAPI channel to ensure blinded analysis. Synaptic structures were identified by co-localisation of Bassoon and PSD95 using a custom-developed ImageJ macro (https://github.com/MRC-LMB-Light-Microscopy-Facility/imagej-macro-cytoplasmic-3d-spot-overlap DOI: 10.5281/zenodo.14562113), with synapse numbers normalised to image volumes.

### Single cell RNA sequencing

For the human and mouse single cell RNA sequencing time course analysis, a total of 48 human and 24 mouse samples were collected at indicated days, with 3 ALI-COs from two independent batches per timepoint. ALI-COs were collected using a cut p1000 tip, transferred to a 15 ml tube pre-coated with BSA and washed once with PBS. Accumax solution (Merck, A7089), 1 ml for human and 200 μl for mouse, was added to the ALI-COs, tissues were gently resuspended 3-5x using a p1000 tip to break it up in smaller pieces, followed by a 15 min incubation at 37°C on a shaker (57 RPM, 25 cm orbit) with gentle mixing every 5 min. The enzymatic activity was stopped by adding 1:10 Fetal Bovine Serum (FBS; Merck, F2442), the cell suspension was gently resuspended another 3-5x and placed on ice to further halt enzymatic reactions. The cell solution was filtered through a 70 μm strainer, centrifuged (300 *g*, 10 min, 4°C) and resuspended in cell prefixation buffer (Parse Biosciences). Sample fixation and library preparation were performed following the Evercode v3 Parse Biosciences protocol, and samples were sequenced using two lanes of the NovaSeq S2.

Mouse and human samples were combined and demultiplexed during subsequent bioinformatic analysis. For species demultiplexing, raw reads were aligned to a combined Human (GRCh38) and Mouse reference (GRCm39) using Split Pipeline v1.0.3 provided by Parse Biosciences. Mapping rates for transcripts to each species reference genome were calculated for every cell based on the obtained counts matrix. Species were demultiplexed by comparing the mapping rates and assigned as the species of the higher mapping rate genome. The cell barcodes of human and mouse assigned cells were extracted and used to subset counts matrices obtained by aligning the raw reads to human and mouse genomes separately. These counts matrices were treated separately through the following analysis steps. For both the human and mouse dataset, as initial quality control, cells were not included if they had: lower than 300 or higher than 15000 UMI counts, fewer than 200 (human) and 300 (mouse) or more than 15000 genes expressed, >10% mitochondrial UMI fractions or >25% ribosomal UMI fractions. Uninformative genes were removed by filtering out genes expressed in less than 1% of the cells. Data was normalised using Scanpy’s `normalize_total` function, and variance was stabilised by applying Scanpy’s `log1p` function (ref). Highly variable genes (HVGs) were identified using Scanpy’s `highly_variable_genes` function (min_mean = 0.05, max_mean = 3, min_disp = 0.75) to select the most informative genes for downstream dimensionality reduction and clustering ^109^. Dimensionality reduction was performed by calculating the first 30 Principal Component Analysis (PCA) based on the HVGs. Uniform Manifold Approximation and Projection (UMAP) was then obtained based on the first 30 PCAs for human or 6 PCAs for mouse, with the specific number determined from elbow plots. The clusters were calculated using Scanpy’s `leiden` function with a 0.25 resolution for human or 0.13 resolution for mouse. Clusters were annotated based on average marker gene expression in the clusters.

In the human dataset, for simplicity of data visualization, two clusters expressing neural progenitor markers were manually merged, as they did not exhibit clear neural progenitor subtype differentiation and were not central to this study. Pseudotime and trajectory analysis was then performed using Palantir ^110^. We found two trajectories and annotated them as ‘Cortical Neurogenesis’ and ‘Choroid Plexus Development’ based on marker gene expression trends for both trajectories along pseudotime. Based on their pseudotime expression profile, genes in the dataset were assigned into 7 different clusters using Palantir’s `cluster_gene_trends` function. The gene clusters were then subjected to GO term enrichment analysis, which informed analysis of differential timing of various stages in neuronal maturation.

In the mouse dataset, for analysis of cortical development, the clusters ‘Radial Glia (late)’, ‘Radial Glia (early) & Intermediate Progenitors’, and ‘Cortical Neurons’ were subsetted from the full dataset (Extended Data Fig 1i). The first 30 PCAs for the subset were calculated, and based on the elbow plot the first 5 PCAs were carried forward to compute the UMAP coordinates. The clusters were calculated using Scanpy’s `leiden` function with resolution of 0.08, and annotated as described before. For the pseudotime analysis of cortical neurogenesis, the clusters ‘Radial Glia (late)’, ‘Radial Glia (early) & Intermediate Progenitors’, ‘Intermediate Progenitors & Cortical Neurons (early)’ and ‘Cortical Neurons (late) clusters were subsetted from the cortical development dataset (Fig 1e). Dimensionality reductions were recalculated as described fore, with the first 6 PCAs carried forward, and a UMAP was calculated with a 0.08 cluster resolution. For simplicity of data visualization, two clusters expressing radial glia markers were manually merged, as they did not exhibit clear neural progenitor subtype differentiation and were not central to this study. Pseudotime and trajectory analysis was then performed using Palantir^110^, revealing a rather small group of 131 cells, representing 1.8% of the total dataset, in the Radial Glia cluster as abnormally mature endpoint. After careful examination these cells turned out to have unexpectedly low counts, as comparing this cell population with the remaining cells in the dataset the Mann-Whitney U revealed a significantly reduced count with a p-value of 3.94e^-39, and compared to other cells in the ‘Radial Glia’ cluster the p-value was 2.83e^-06. The cells were removed from the dataset and the pseudotime was calculated showing two clear trajectories, which were annotated as ‘Cortical neurogenesis’ and ‘Cortical gliogenesis’ based on the marker genes along the pseudotimes trajectories. Genes in the cortical neurogenesis trajectory were then assigned into 8 different clusters based on their expression dynamics, with clusters being subjected to GO term enrichment analysis, as described above.

To perform pseudotime alignment between mouse and human portion of the data, the final subsets were renormalised together and the gene space has been subsetted to the common overlap. Trajectories were aligned using Genes2Genes method (bin = 14) using human part of the data as reference and mouse part as query ^111^. The average alignment based on all the shared genes, had then been used to subset the aligned sections of pseudotimes in both human and mouse dataset. Gene trends along the pseudotime sections were recalculated using `palantir.presults.compute_gene_trends()` (expression_key=“MAGIC_imputed_data”) and plotted individually or averaged per gene module.

For the single cell RNA sequencing analysis of human organoids treated with DMSO (control) or BayK8644 (L-Type VGCC agonist), we used human brain organoid slices cultured in micropatterned devices and subjected to pharmacological treatment, all using similar protocols as described above, that were snapfrozen at day 90 and stored at -80C. For each condition, we performed 2 reactions using samples from independent batches, with 2 organoids pooled for each reaction. Samples were homogenized in 1 ml NP-40 0.1 lysis buffer (10 mM Tris-HCl pH 7.4, 10 mM NaCl, 3 mM MgCl2, 5 mM CaCl2, 0.1% (v/v) IGE-PAL CA-630; supplemented with 1 mM DTT, 1x protease inhibitor cocktail, EDTA-free and 0.1 U/µl RNase inhibitor (Sigma, 03335399001)), by 10 strokes with pestle A and 7 strokes with pestle B. Cells were filtered through a 20 um strainer (PluriSelect, 43-50020-01), washed with 1 ml nuclei isolation wash buffer (10 mM Tris-HCl pH 7.4, 10 mM NaCl, 3 mM MgCl2, 0.1% (v/v) Tween-20; supplemented with 1 mM DTT, 1x 1protease inhibitor cocktail, EDTA-free and 0.1 U/µl RNase inhibitor (Sigma, 03335399001)), pelleted (700 *g*, 5 min, 4°C), resuspended in 1 ml nuclei isolation wash buffer. Cells were pelleted again (700 *g*, 5 min, 4°C), resuspended in 100 µl multiome lysis buffer (10 mM Tris-HCl pH 7.4, 1% BSA, 10 mM NaCl, 3 mM MgCl2, 0.01% (v/v) Tween-20%, 0.01% (v/v) IGE-PAL CA-630, 0.001% (w/v) Digitonin; supplemented with 1 mM DTT, 1x 1protease inhibitor cocktail, EDTA-free and 0.1 U/µl RNase inhibitor (Sigma, 03335399001)) by gently pipetting 5x, incubated for 2 min on ice. Then 500 ul nuclei isolation wash buffer was added, cells were pelleted (700 *g*, 5 min, 4°C) and resuspended in Diluted Nuclei Buffer (10xgenomics, 2000207). Samples were processed following the Chromium Next GEM Single Cell Multiome ATAC + Gene Expression user guide (10xgenomics). Cells were sequenced on a lane of the NovaSeq X (10B), and only the scRNA-seq data, but not the sc-ATAC data, was carried forward for analysis, due to a technical issue leading to incomplete index sequences of the ATAC reads.

As initial quality control, the dataset was filtered to exclude cells in the population (normalized per sample): <5% total genes expressed, <5% UMI counts, >99.5% UMI counts, >95% exhibiting the highest mitochondria fractions. Data was normalised and log-transformed using the Scanpy functions mentioned above. The top 4000 HVGs per sample were identified using Scanpy, as described above, for which uninformative genes (<0.1% expression in cells or expressed in <10 cells) were removed. UMAP was then obtained based on the first 11 PCAs (cosine, k=15, min_dist=0.8), as determined from an elbow plot, followed by harmony integration using Scanpy’s external function `harmony_integrate` (theta=2.0). The clusters were computed with a resolution of 0.07 and annotated by analysing marker gene expression, as described above.

### Statistical analysis

Statistical details are provided in the corresponding figure legends. *P*-values are annotated as followed: ns P > 0.05, \**P* < 0.05, \*\**P* < 0.01, \*\*\**P* < 0.001 and \*\*\*\**P* < 0.0001. Data processing and statistical analyses were carried out in GraphPad Prism (version 10.0) or in Jupyter Notebooks (version 4.2.3) with Python for scRNA-seq analysis.

## ACKNOWLEDGEMENTS

The authors would like to thank members of the Lancaster lab for helpful feedback and discussions. We also thank Jess L. Sevetson and Alex Justin for their support with bioengineering the micropatterned devices. Thanks to Marta Shahbazi for providing the mouse E14 cell line and to Casper Hoogenraad for gifting the MARCKS-GFP plasmid. This work was supported by the Medical Research Council (MC_UP_1201/9) and the European Research Council (ERC-STG, 757710) to MAL. FWL is a non-stipendiary European Molecular Biology Organisation Postdoctoral Fellow (EMBO, ALTF 845-2021) and supported by the Netherlands Organisation for Scientific Research (NWO-Rubicon, 019.211EN.032ß).

## AUTHOR CONTRIBUTIONS

FWL and MAL designed the study and wrote the manuscript. FWL performed experiments and analysed data. HMS generated and analysed the cross-species synaptogenesis data, under supervision of FWL, and conducted bioinformatic analysis. II-R and LG performed initial bioinformatic analysis. KV, MM and JM generated master molds for micropatterned devices, under supervision of MT. NZG conducted and analysed the gliogenesis data, under supervision of FWL. JB and US developed methods for image analysis. DJLDS and MAL generated mouse spinal cord explant tissues. DJLDS, MAL, IC and LP contributed to sample collection for RNA sequencing.

---

**Extended Data Figure 1.**
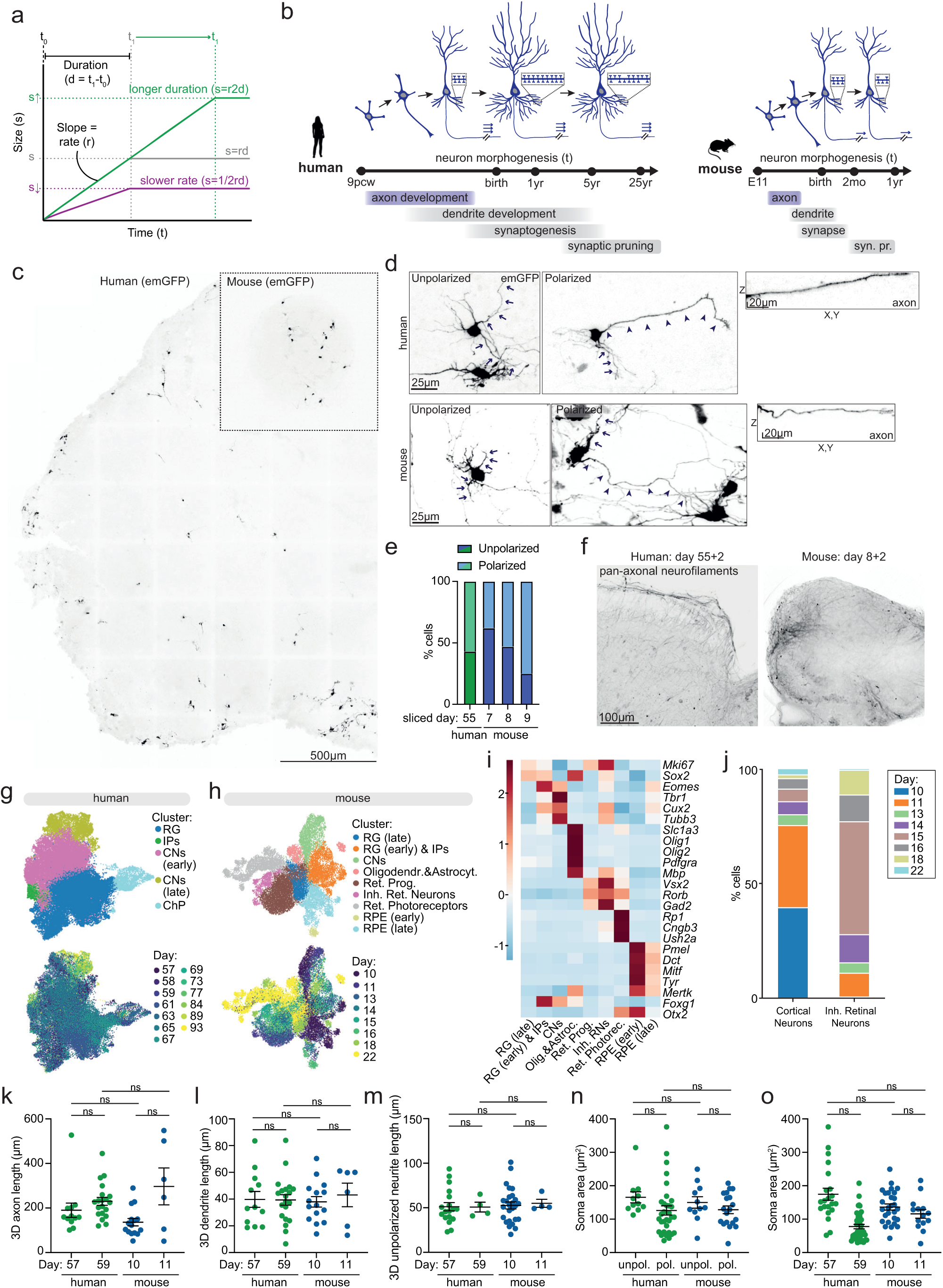
a. Hypothetical model showing the relationship between morphological size (s), developmental rate (r) and developmental duration (d). b. Schematic illustration of the sequential developmental steps and timeline of human and mouse cortical neuron morphogenesis, with species-specific neuron morphologies drawn to scale^17,24,25,27–29,45,112,113^. c. Representative image of a day 69 human and a day 15 mouse ALI-CO with sparsely labelled emGFP-positive cells. d. Representative images of sparsely labelled emGFP-positive unpolarized neurons (arrowheads marking unpolarised neurites) and polarized neurons (arrowheads mark axons, arrows mark longest dendrite) in human and mouse ALI-COs. Zoom shows a line plot of the axons in Z. e. Quantifications of the relative proportion of sparsely labelled emGFP-positive unpolarized and polarized neurons 24-36 hours after slicing in human ALI-COs, sliced at day 55, and mouse ALI-COs, sliced at day 7, 8 or 9. Hm day 55: n=21 neurons, Ms day 7: n=22, Ms day 8: n=51, Ms day 9: n=8. f. Similar axon morphologies at stage-matched timepoints in human (sliced at day 55, analysed 2 and 42 days later) and mouse (sliced at day 8, analysed 2 and 14 days later) ALI-COs immunostained for pan-axonal neurofilaments. g. UMAP embedding of the human brain organoid dataset. Cells are colored cluster identity or timepoints. h. UMAP embedding of the mouse brain organoid dataset. Cells are colored cluster identity or timepoints. i. Heatmap of representative marker genes of clusters in (g). j. Bar plot of timepoints representation in the cortical neuron and inhibitory retinal neuron clusters in (i), indicating that only cortical neurons are present at day 10. k. Quantifications of 3D axon length in sparsely labelled emGFP-positive polarised neurons on day 57 and 79 in human ALI-COs, and day 10 and 11 in mouse ALI-COs. Hm day 57: n=12 neurons, Hm day 59: n=20, Ms day 10: n= 42, Ms day 11: n=6. l. Quantifications of 3D length of the longest dendrite in sparsely labelled emGFP-positive polarised neurons on day 57 and 79 in human ALI-COs, and day 10 and 11 in mouse ALI-COs. Hm day 57: n=12 neurons, Hm day 59: n=20, Ms day 10: n= 42, Ms day 11: n=6. m. Quantifications of 3D unpolarized neurite length in sparsely labelled emGFP-positive unpolarised neurons on day 57 and 79 in human ALI-COs, and day 10 and 11 in mouse ALI-COs. Hm day 57: n=18 neurons, Hm day 59: n=4, Ms day 10: n= 74, Ms day 11: n=4. n. Quantifications of soma areas in sparsely labelled emGFP-positive unpolarised and polarised neurons in human (day 57-59) and mouse (day 10-11) ALI-COs. Hm day 57: n=21 neurons, Hm day 59: n=40, Ms day 10: n= 57, Ms day 11: n=14, Ms day 15: n=7. p. Quantifications of soma areas in sparsely labelled emGFP-positive unpolarised and polarised neurons on day 57 and 79 in human ALI-COs, and day 10 and 11 in mouse ALI-COs. Hm unpolarized: n=11 neurons, Hm polarized: n=32, Ms unpolarized: n=39, Ms polarized: n=48. Syn., synaptic. emGFP, emerald green fluorescent protein. Unpol., unpolarized. RG., radial glia. IPs., intermediate progenitors. CNs, cortical neurons. ChP, choroid plexus. Oligodendr., oligodendrocytes. Astrocyt., astrocytes. Ret., retinal. Prog., progenitors. Inh., inhibitory. RPE, retinal pigmented epithelium. All data are shown as mean ± SEM; ns *P* > 0.05 as determined by unpaired *t* tests.

**Extended Data Figure 2.**
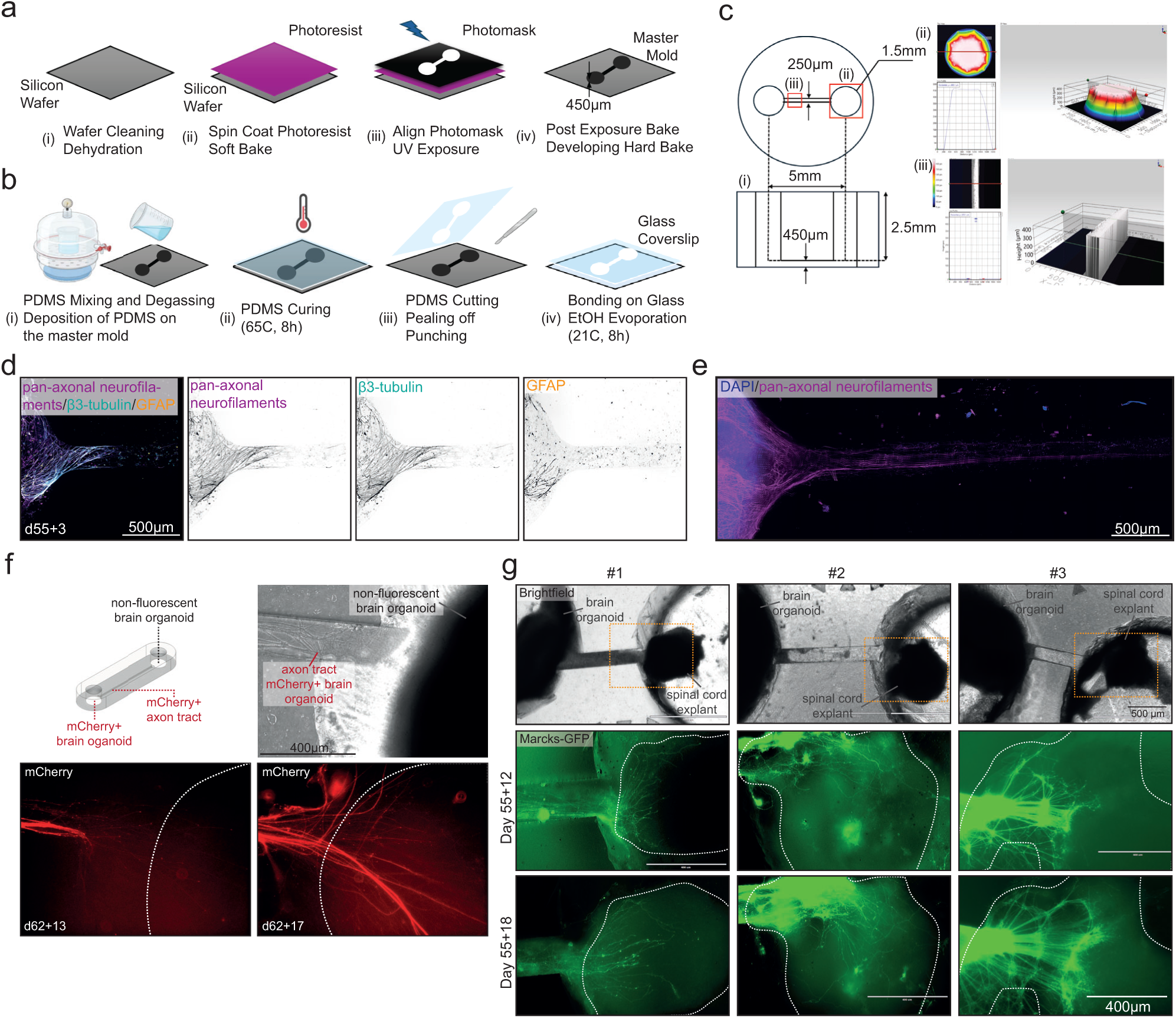
a. Photolithography process for master mold fabrication. (i) Wafer cleaning (15 min, Nanostrips® and 15min, DI water), dehydration (15 min, 150°C). (ii) Spin coat SU8-100 photoresist (Step 1: 500 rpm at 100 rpm/sec, 10 sec. Step 2: 1000 rpm at 300 rpm/sec, 30 sec), soft bake photoresist (Step 1: 30 min, 65°C. Step 2: 90 min, 95°C). (iii) Prepare Mylar photomask (Step 1: CAD design and conversion. Step 2: order printing the CAD design), UV exposure at OAI Model 204 Mask Aligner (Step 1: align photomask and substrate. Step 2: UV exposure time 115.21 sec, channel intensity (mW/cm^2^):7.2, exposure energy (mJ/cm^2^): 829.5). iv. Post exposure bake (Step1: 1min, 65°C. Step 2: 20 min, 95°C), developing (Step 1: 20 min SU8 developer. IPA rinse, nitrogen dry. Step 2: hard bake, 15 min, 150°C). b. Soft lithography process for PDMS microstructure fabrication. (i) PDMS mixing (10:1), degassing, and deposition onto fabricated master mold. (ii) PDMS curing (65°C for 8 hours). (iii) Cutting and peeling off the PDMS, punching the milli wells. (iv) Oxygen plasma exposure (30 sec) for bonding devices onto glass and placing onto a hot plate (80°C) for 45 min. c. Device features. (i) 2D schematic of the microstructure master mold in top and side view with dimensions. (ii) Thickness characterization of the milli well. (iii) Thickness characterization of the master mold microchannel. d. Human brain organoid slice cultures placed in the chambers at day 55, fixed 3 days later and immunostained for pan-axonal filaments, β3-tubulin and GFAP. e. Human brain organoid slice cultures placed in the chambers at day 55, fixed 14 days later and immunostained for pan-axonal filaments and DAPI. f. Schematic illustration of experimental set-up for the target tissue experiment. RFP-positive axons from an RFP-positive human brain organoid slice culture in a short micropatterned device channel extending on to another non-fluorescent human organoid, outline indicated by dashed line, 12 days and 18 days after being placed in the chamber (at day 62). g. Brightfield images of micropatterned devices with a human organoid slice culture in the left chamber and a mouse spinal cord explant in the right chamber. Zoom shows Marcks-GFP-positive axons from a human brain organoid slice culture extending onto a mouse spinal cord explant, indicated by dashed line, 12 days and 18 days after being placed in the chamber (at day 55). Hm, human. Ms, mouse.

**Extended Data Figure 3.**
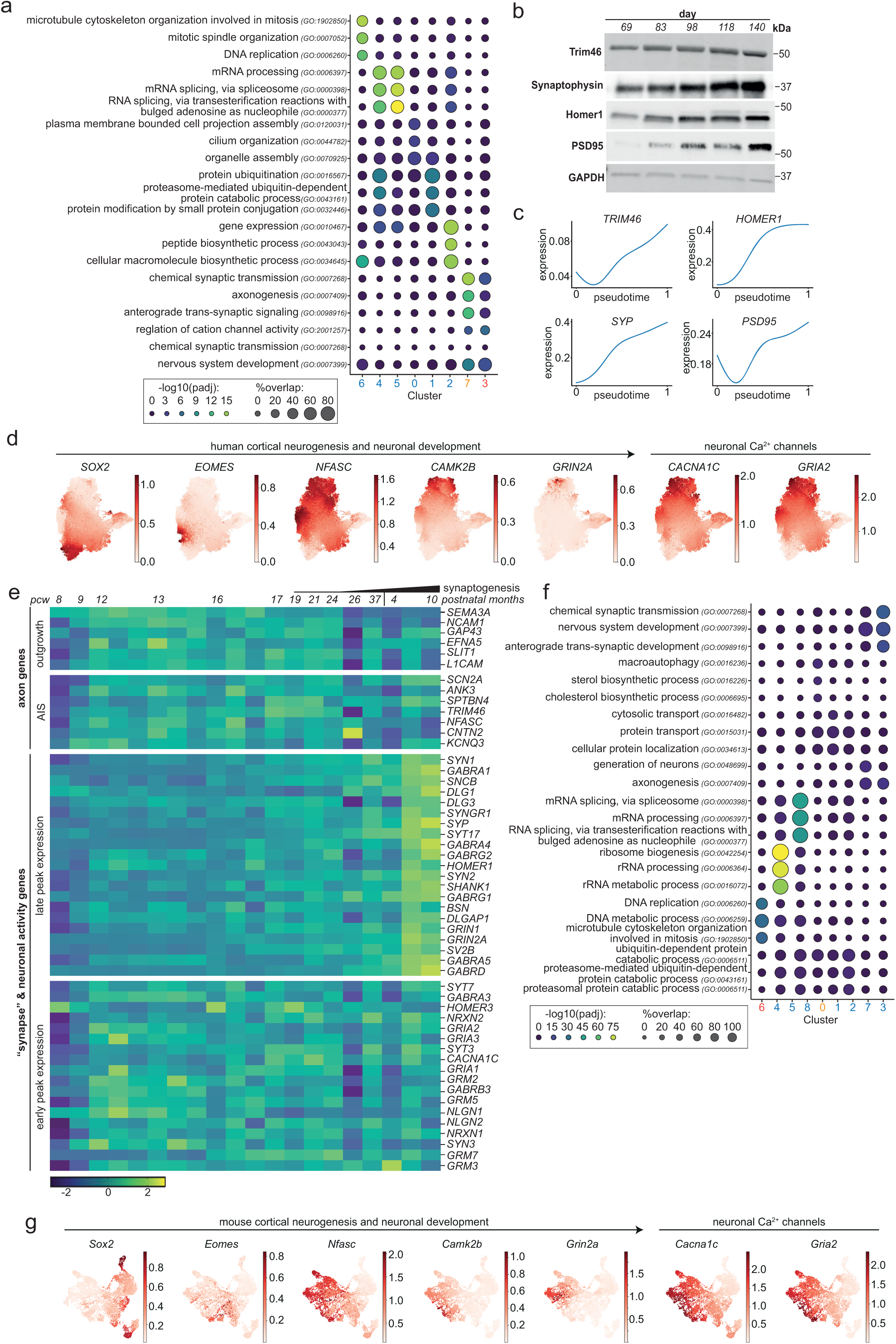
a. Dotplot showing top 3 enriched GO terms for each gene cluster in (Fig 2b). b. Western blot of human ALI-COs at different timepoints showing temporal expression profiles of neuron maturation proteins. c. Human pseudotemporal scRNA-seq expression patterns of neuronal maturation genes (same as in b). d. Feature plots of the human scRNA-seq dataset showing expression profiles of representative cortical neurogenesis, neuronal development and neuronal calcium marker genes. e. Heatmap of RNA expression data (Z scores of log2 RPKM values) from the human prefrontal cortex *in vivo* (data from Brainspan) covering embryonic and early postnatal timepoints. f. Dotplot showing top 3 enriched GO terms for each gene cluster in (Fig 2j) g. Feature plots of the human scRNA-seq dataset showing expression profiles of representative cortical neurogenesis, neuronal development and neuronal calcium marker genes. Pcw, postconception week. AIS, axon initial segment.

**Extended Data Figure 4.**
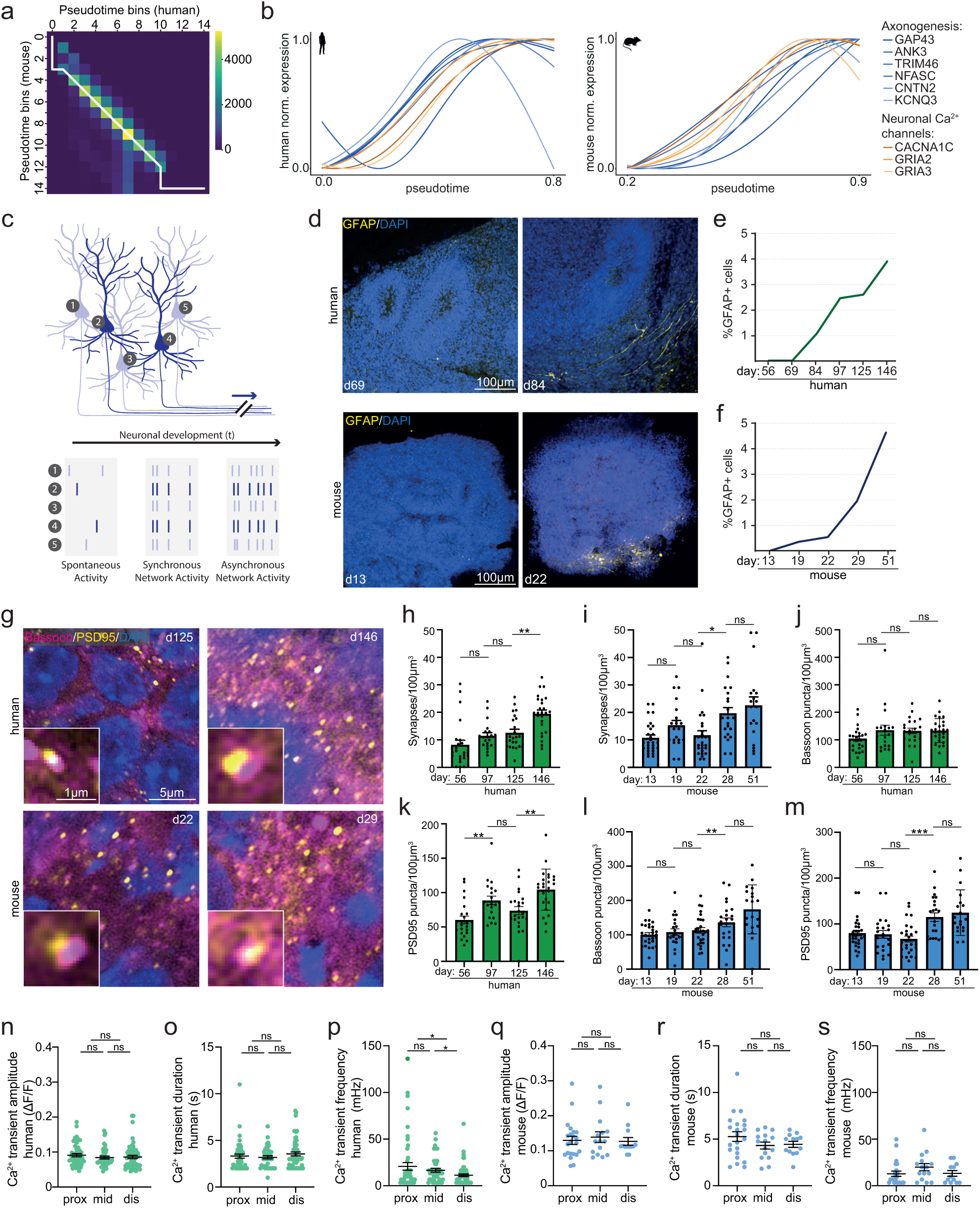
a. Heatmap showing alignment of 15 pseudotime bins in human and mouse. b. Expression profile of axonogenesis and neuronal calcium channel genes along aligned pseudotime trajectories in human and mouse. c. Schematic of different modes of neural activity during neuron development ^74–76^. d. Representative images of cryosections of human and mouse ALI-COs at indicated days immunostained for GFAP and DAPI. e. Quantifications of proportion of GFAP-positive cells in human ALI-COs across different timepoints. Day 56: n=14 regions of interest, Day 69: n=14, Day 84: n= 14, Day 97: n=13, Day 125: n=14, Day 14: n=13. f. Quantifications of proportion of GFAP-positive cells in mouse ALI-COs across different timepoints. Day 13: n=14 regions of interest, Day 19: n=14, Day 22: n=14, Day 19: n=14, Day 42: n=9, Day 51: n=13. g. Representative images of cryosections of human and mouse ALI-COs immunostained for bassoon, PSD95 and DAPI, showing an increase of synaptic puncta at indicated timepoints. Zoom represents a co-localisation of pre- and postsynaptic structure. h. Quantifications of number of co-localising bassoon and PSD95 puncta in cryosections of human ALI-COs at different timepoints. Day 56: n=24 regions of interest, Day 69: n=13, Day 84: n=11, Day 97: n=19, Day 125: n=22, Day 147: n=14. i. Quantifications of number of co-localising bassoon and PSD95 puncta in cryosections of mouse ALI-COs at different timepoints. Day 9: n=10 regions of interest, Day 13: n=27, Day 19: n=21, Day 22: n=27, Day 29: n=21, Day 51: n=17. j. Quantifications of number of Bassoon puncta in cryosections of human ALI-COs at different timepoints. Day 56: n=24 regions of interest, Day 69: n=13, Day 84: n=11, Day 97: n=19, Day 125: n=22, Day 147: n=14. k. Quantifications of number of bassoon puncta in cryosections of mouse ALI-COs at different timepoints. Day 9: n=10 regions of interest, Day 13: n=27, Day 19: n=21, Day 22: n=27, Day 29: n=21, Day 51: n=17. l. Quantifications of number of PSD95 puncta in cryosections of human ALI-COs at different timepoints. Day 56: n=24 regions of interest, Day 69: n=13, Day 84: n=11, Day 97: n=19, Day 125: n=22, Day 147: n=14. m. Quantifications of number of PSD95 puncta in cryosections of mouse ALI-COs at different timepoints. Day 9: n=10 regions of interest, Day 13: n=27, Day 19: n=21, Day 22: n=27, Day 29: n=21, Day 51: n=17. n. Quantifications of average mean amplitudes of spontaneous calcium transients in proximal, mid or distal regions of axon tracts of human (day 69) organoid slice cultures in micropatterned devices expressing GCaMP7f. Proximal: n=50 axons, mid: n=43, distal n=58. o. Quantifications of average event durations of spontaneous calcium transients in proximal, mid or distal regions of axon tracts of human (day 69) organoid slice cultures in micropatterned devices expressing GCaMP7f. Proximal: n=50 axons, mid: n=43, distal n=58. p. Quantifications of average frequencies of spontaneous calcium transients in proximal, mid or distal regions of axon tracts of human (day 69) organoid slice cultures in micropatterned devices expressing GCaMP7f. Proximal: n=50 axons, mid: n=43, distal n=58. q. Quantifications of average mean amplitudes of spontaneous calcium transients in proximal, mid or distal regions of axon tracts of mouse (day 15) organoid slice cultures in micropatterned devices expressing GCaMP7f. Proximal: n=23 axons, mid: n=15, distal n=13. r. Quantifications of average event durations of spontaneous calcium transients in proximal, mid or distal regions of axon tracts of mouse (day 15) organoid slice cultures in micropatterned devices expressing GCaMP7f. Proximal: n=23 axons, mid: n=15, distal n=13. s. Quantifications of average frequencies of spontaneous calcium transients in proximal, mid or distal regions of axon tracts of mouse (day 15) organoid slice cultures in micropatterned devices expressing GCaMP7f. Proximal: n=23 axons, mid: n=15, distal n=13. Prox, proximal. Dis, distal. Norm., normalized. All data are shown as mean ± SEM; ns *P* > 0.05, \**P* < 0.05, \*\**P* < 0.1, \*\*\**P* < 0.001 as determined by unpaired *t* tests.

**Extended Data Figure 5.**
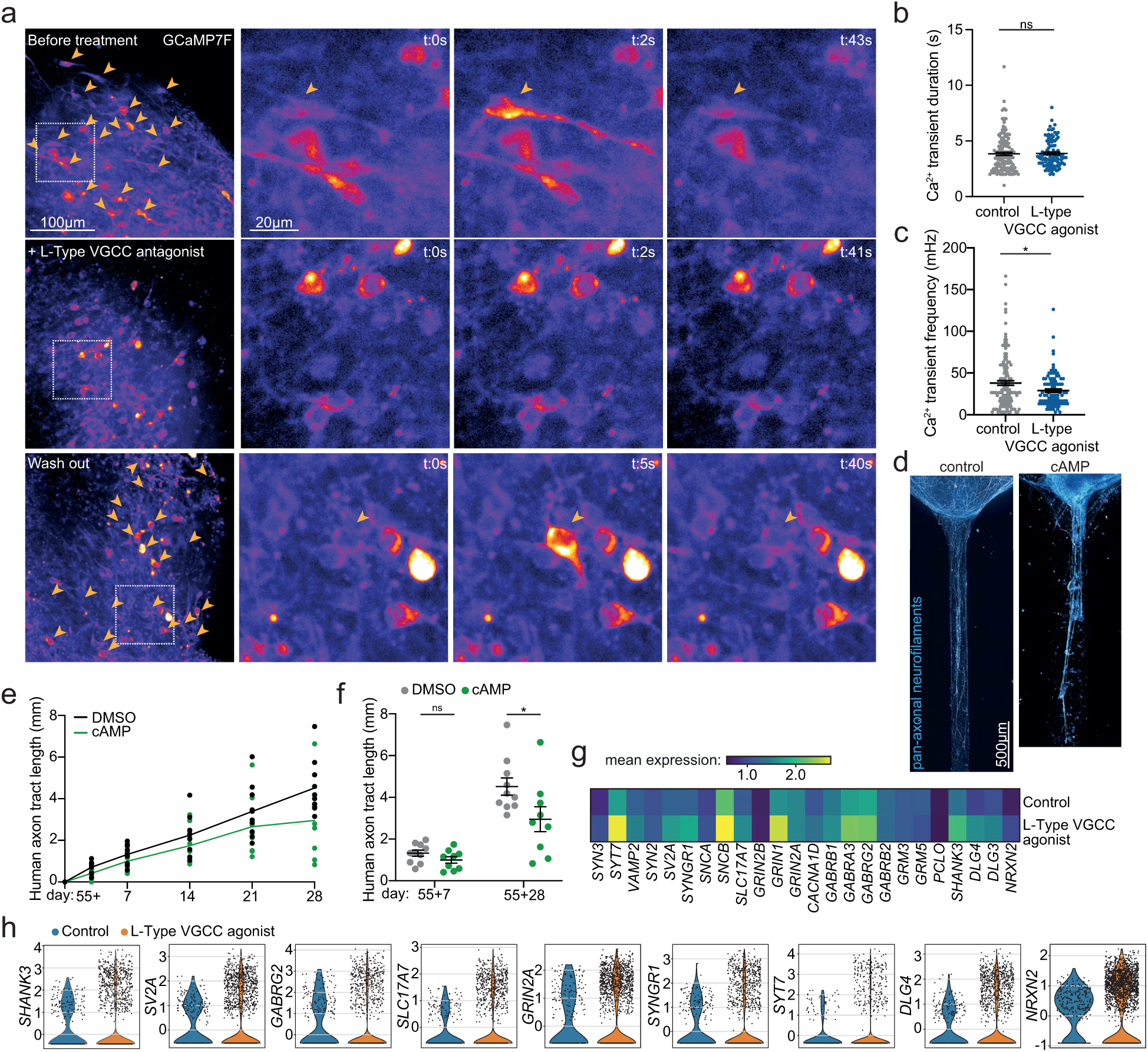
a. Human (day 96) brain organoid slices in micropatterned devices expressing GCaMP7f before treatment, after 1 hour treatment with nimodipine (L-Type VGCC antagonist) and 3 hours after wash-out of treatment. Arrowheads mark axons showing calcium activity in a five minute timelapse, indicating nearly complete perturbation of calcium dynamics upon L-Type VGCC blocking and full recovery after wash-out. Zooms show stills of representative timelapse of individual neurons (marked by arrowheads). b. Quantifications of average frequencies of spontaneous calcium transients in human brain organoid slice cultures (day 70) in micropatterned devices expressing GCaMP7f and treated with DMSO (control) or BayK8644 (L-Type VGCC agonist). Control: n=137 axons, L-Type VGCC agonist: n=100. c. Quantifications of average event durations of spontaneous calcium transients in human brain organoid slice cultures (day 70) in micropatterned devices expressing GCaMP7f and treated with DMSO (control) or BayK8644 (L-Type VGCC agonist). Control: n=137 axons, L-Type VGCC agonist: n=100. d. Representative images of axon tracts of human (day 90) brain organoid slices in micropatterned devices immunostained for pan-axonal neurofilaments and treated with DMSO (control) or cAMP. e. Quantifications of axon tract length of human brain organoid slice cultures in micropatterned devices, treated with DMSO (control) or cAMP, at different timepoints. n=10 organoids. Mixed-effects model with Geisser-Greenhouse correction P<0.0001 for time, p=0.0766 for treatments (based on average length across timepoints). f. Quantifications of axon tract length of human brain organoid slice cultures in micropatterned devices, treated with DMSO (control) or cAMP, at day 55+7 and 55+28. Control: n=10 organoids (55+7) and n=10 (55+28), cAMP: n=9 (55+7) and n=9 (55+28). Multiple unpaired t-test with two-stage step-up (Denjamini, Krieger and Yekutieli; FDR 10%). g. Heatmap of synapse gene expression of non-zero expressing cells in deep layer neuron cluster in (Fig 4h), comparing control and L-type VGCC agonist conditions. h. Violin plots of synapse gene expression of cells in upper layer neuron cluster in (Fig 4h), comparing control and L-type VGCC agonist conditions. All data are shown as mean ± SEM; ns *P/Q* > 0.05, \**P* < 0.05, as determined by unpaired *t* tests unless stated differently.

**Supplementary Table 1.** Clustering of genes from the human scRNA-seq dataset based on their distinct expression profiles along the cortical neurogenesis pseudotime trajectory, with analysis of synapse and neural activity genes in clusters 3 and 7.

**Supplementary Table 2.** Clustering of genes from the mouse scRNA-seq dataset based on their distinct expression profiles along the cortical neurogenesis pseudotime trajectory, with analysis of synapse and neural activity genes in clusters 0 and 6.

**Supplementary Video 1.**

Timelapse of spontaneous calcium transients in an axon tract of a human (day 69) brain organoid slice in micropatterned devices expressing GCaMP7f.

**Supplementary Video 2.**

Timelapse of spontaneous calcium transients in an axon tract of a mouse (day 15) brain organoid slice in micropatterned devices expressing GCaMP7f.

**Supplementary Video 3.**

Timelapse of synchronous calcium bursts in an axon tract of a human (day 82) brain organoid slice in micropatterned devices expressing GCaMP7f.

**Supplementary Video 4.**

Timelapse of synchronous calcium bursts in an axon tract of a mouse (day 19) brain organoid slice in micropatterned devices expressing GCaMP7f.

**Supplementary Video 5.**

Timelapse of spontaneous calcium transients in an axon tract of a human (day 69) brain organoid slice in micropatterned devices expressing GCaMP7f and treated with DMSO.

**Supplementary Video 6.**

Timelapse of spontaneous calcium transients in an axon tract of a human (day 69) brain organoid slice in micropatterned devices expressing GCaMP7f and treated with nimodipine (L-Type VGCC antagonist).

**Supplementary Video 7.**

Timelapse of spontaneous calcium transients in an axon tract of a human (day 69) brain organoid slice in micropatterned devices expressing GCaMP7f and treated with BayK8644 (L-Type VGCC agonist).

**Supplementary Video 8.**

Timelapse of calcium activity in a human brain organoid slice (day 96) expressing GCaMP7f before treatment.

**Supplementary Video 9.**

Timelapse of calcium activity in a human brain organoid slice (day 96) expressing GCaMP7f after treatment with nimodipine (L-Type VGCC antagonist).

**Supplementary Video 10.**

Timelapse of calcium activity in a human brain organoid slice (day 96) expressing GCaMP7f after washout upon treatment with nimodipine (L-Type VGCC antagonist).

## REFERENCES

1. Venditti, C., Baker, J. & Barton, R. A. Co-evolutionary dynamics of mammalian brain and body size. *Nat*. Ecol. Evol. 8, 1534–1542 (2024).

2. Masoli, S. et al. Human outperform mouse Purkinje cells in dendritic complexity and computational capacity. (2023) doi:10.1101/2023.03.08.531672.

3. Uzquiano, A. & Arlotta, P. Brain organoids: the quest to decipher human-specific features of brain development. Curr Opin Genet Dev 75, 101955 (2022).

4. DeCasien, A. R., Barton, R. A. & Higham, J. P. Understanding the human brain: insights from comparative biology. Trends Cogn. Sci. 26, 432–445 (2022).

5. Libé-Philippot, B. & Vanderhaeghen, P. Cellular and Molecular Mechanisms Linking Human Cortical Development and Evolution. Annu Rev Genet 55, 1–27 (2021).

6. Smaers, J. B. et al. The evolution of mammalian brain size. Sci. Adv. 7, eabe2101 (2021).

7. Beaudet, A., Du, A. & Wood, B. Evolution of the modern human brain. Prog Brain Res 250, 219–250 (2019).

8. Dicke, U. & Roth, G. Neuronal factors determining high intelligence. Philosophical Transactions Royal Soc B Biological Sci 371, 20150180 (2016).

9. Mohan, H. et al. Dendritic and Axonal Architecture of Individual Pyramidal Neurons across Layers of Adult Human Neocortex. Cereb Cortex 25, 4839–4853 (2015).

10. Bianchi, S. et al. Dendritic Morphology of Pyramidal Neurons in the Chimpanzee Neocortex: Regional Specializations and Comparison to Humans. Cereb Cortex 23, 2429–2436 (2013).

11. DeFelipe, J. The Evolution of the Brain, the Human Nature of Cortical Circuits, and Intellectual Creativity. Front Neuroanat 5, 29 (2011).

12. Azevedo, F. A. C. et al. Equal numbers of neuronal and nonneuronal cells make the human brain an isometrically scaled-up primate brain. J. Comp. Neurol. 513, 532–541 (2009).

13. Herculano-Houzel, S., Collins, C. E., Wong, P. & Kaas, J. H. Cellular scaling rules for primate brains. Proc National Acad Sci 104, 3562–3567 (2007).

14. Elston, G. N. et al. Specializations of the granular prefrontal cortex of primates: Implications for cognitive processing. Anatomical Rec Part Discov Mol Cell Evol Biology 288A, 26–35 (2006).

15. Elston, G. N., Benavides-Piccione, R. & DeFelipe, J. A Study of Pyramidal Cell Structure in the Cingulate Cortex of the Macaque Monkey with Comparative Notes on Inferotemporal and Primary Visual Cortex. Cereb Cortex 15, 64–73 (2005).

16. Elston, G. N., Benavides-Piccione, R. & DeFelipe, J. The Pyramidal Cell in Cognition: A Comparative Study in Human and Monkey. J Neurosci 21, RC163–RC163 (2001).

17. Jacobs, B. et al. Regional Dendritic and Spine Variation in Human Cerebral Cortex: a Quantitative Golgi Study. Cereb Cortex 11, 558–571 (2001).

18. Zhang, K. & Sejnowski, T. J. A universal scaling law between gray matter and white matter of cerebral cortex. Proc National Acad Sci 97, 5621–5626 (2000).

19. Wei, Y. et al. Cross-Species Cortical Geometry Reveals Conserved Gradients Across Primates and Human-Specific Expansion. bioRxiv 2025.11.28.691123 (2025) doi:10.1101/2025.11.28.691123.

20. Donahue, C. J., Glasser, M. F., Preuss, T. M., Rilling, J. K. & Essen, D. C. V. A Quantitative Assessment of Prefrontal Cortex in Humans Relative to Nonhuman Primates. Biorxiv 233346 (2017) doi:10.1101/233346.

21. Hill, J. et al. Similar patterns of cortical expansion during human development and evolution. Proc National Acad Sci 107, 13135–13140 (2010).

22. Schoenemann, P. T., Sheehan, M. J. & Glotzer, L. D. Prefrontal white matter volume is disproportionately larger in humans than in other primates. Nat Neurosci 8, 242–252 (2005).

23. Lindhout, F. W., Krienen, F. M., Pollard, K. S. & Lancaster, M. A. A molecular and cellular perspective on human brain evolution and tempo. Nature 630, 596–608 (2024).

24. Wildenberg, G., Li, H. & Kasthuri, N. The Development of Synapses in Mouse and Macaque Primary Sensory Cortices. Biorxiv 2023.02.15.528564 (2023) doi:10.1101/2023.02.15.528564.

25. Zeiss, C. J. Comparative Milestones in Rodent and Human Postnatal Central Nervous System Development. Toxicol Pathol 49, 1368–1373 (2021).

26. Meng, Y., Jiang, J., Bachevalier, J., Zhang, X. & Chan, A. W. S. Developmental Whole Brain White Matter Alterations in Transgenic Huntington’s Disease Monkey. Sci Rep-uk 7, 379 (2017).

27. Sakai, T. et al. Developmental trajectory of the corpus callosum from infancy to the juvenile stage: Comparative MRI between chimpanzees and humans. Plos One 12, e0179624 (2017).

28. Semple, B. D., Blomgren, K., Gimlin, K., Ferriero, D. M. & Noble-Haeusslein, L. J. Brain development in rodents and humans: Identifying benchmarks of maturation and vulnerability to injury across species. Prog. Neurobiol. 106, 1–16 (2013).

29. Huttenlocher, P. R. & Dabholkar, A. S. Regional differences in synaptogenesis in human cerebral cortex. J Comp Neurol 387, 167–178 (1997).

30. Bourgeois, J. & Rakic, P. Changes of synaptic density in the primary visual cortex of the macaque monkey from fetal to adult stage. J Neurosci 13, 2801–2820 (1993).

31. Rakic, P., Bourgeois, J.-P., Eckenhoff, M. F., Zecevic, N. & Goldman-Rakic, P. S. Concurrent Overproduction of Synapses in Diverse Regions of the Primate Cerebral Cortex. Science 232, 232–235 (1986).

32. Linaro, D. et al. Xenotransplanted Human Cortical *Neuron*s Reveal Species-Specific Development and Functional Integration into Mouse Visual Circuits. Neuron 104, 972–986.e6 (2019).

33. Schörnig, M. et al. Comparison of induced neurons reveals slower structural and functional maturation in humans than in apes. Elife 10, e59323 (2021).

34. Otani, T., Marchetto, M. C., Gage, F. H., Simons, B. D. & Livesey, F. J. 2D and 3D Stem Cell Models of Primate Cortical Development Identify Species-Specific Differences in Progenitor Behavior Contributing to Brain Size. Cell Stem Cell 18, 467–480 (2016).

35. Diaz-Cuadros, M. et al. Metabolic regulation of species-specific developmental rates. Nature 613, 550–557 (2023).

36. Diaz-Cuadros, M. et al. In vitro characterization of the human segmentation clock. Nature 580, 113–118 (2020).

37. Matsuda, M. et al. Species-specific segmentation clock periods are due to differential biochemical reaction speeds. Science 369, 1450–1455 (2020).

38. Rayon, T. et al. Species-specific pace of development is associated with differences in protein stability. Science 369, eaba7667 (2020).

39. Iwata, R. et al. Mitochondria metabolism sets the species-specific tempo of neuronal development. Science 379, eabn4705 (2023).

40. Ciceri, G. et al. An epigenetic barrier sets the timing of human neuronal maturation. Nature 1–10 (2024) doi:10.1038/s41586-023-06984-8.

41. Benito-Kwiecinski, S. et al. An early cell shape transition drives evolutionary expansion of the human forebrain. Cell 184, 2084–2102.e19 (2021).

42. Stepien, B. K., Vaid, S. & Huttner, W. B. Length of the Neurogenic Period—A Key Determinant for the Generation of Upper-Layer Neurons During Neocortex Development and Evolution. Front. Cell Dev. Biol. 9, 676911 (2021).

43. Kanton, S. et al. Organoid single-cell genomic atlas uncovers human-specific features of brain development. Nature 574, 418–422 (2019).

44. Silbereis, J. C., Pochareddy, S., Zhu, Y., Li, M. & Sestan, N. The Cellular and Molecular Landscapes of the Developing Human Central Nervous System. Neuron 89, 248–268 (2016).

45. Haubensak, W., Attardo, A., Denk, W. & Huttner, W. B. Neurons arise in the basal neuroepithelium of the early mammalian telencephalon: A major site of neurogenesis. Proc. Natl. Acad. Sci. 101, 3196–3201 (2004).

46. Donahue, C. J., Glasser, M. F., Preuss, T. M., Rilling, J. K. & Essen, D. C. V. Quantitative assessment of prefrontal cortex in humans relative to nonhuman primates. Proc. Natl. Acad. Sci. 115, E5183–E5192 (2018).

47. Sánchez, D. J. L.-D., Lindhout, F. W., Anderson, A. J., Pellegrini, L. & Lancaster, M. A. Mouse brain organoids model in vivo neurodevelopment and function and capture differences to human. bioRxiv 2024.12.21.629881 (2024) doi:10.1101/2024.12.21.629881.

48. Giandomenico, S. L. et al. Cerebral organoids at the air-liquid interface generate diverse nerve tracts with functional output. Nat Neurosci 22, 669–679 (2019).

49. Paşca, A. M. et al. Functional cortical neurons and astrocytes from human pluripotent stem cells in 3D culture. Nat. Methods 12, 671–678 (2015).

50. Dotti, C., Sullivan, C. & Banker, G. The establishment of polarity by hippocampal neurons in culture. J. Neurosci. 8, 1454–1468 (1988).

51. Lindhout, F. W. et al. Quantitative mapping of transcriptome and proteome dynamics during polarization of human iPSC-derived neurons. Elife 9, e58124 (2020).

52. Chiaradia, I. et al. Tissue morphology influences the temporal program of human brain organoid development. Cell Stem Cell 30, 1351–1367.e10 (2023).

53. Lancaster, M. A. et al. Cerebral organoids model human brain development and microcephaly. Nature 501, 373–379 (2013).

54. Medina-Cano, D. et al. Rapid and robust directed differentiation of mouse epiblast stem cells into definitive endoderm and forebrain organoids. Development 149, (2022).

55. Sarnat, H. B. Axonal pathfinding during the development of the nervous system. Ann. Child Neurol. Soc. 1, 13–23 (2023).

56. McCormick, L. E. & Gupton, S. L. Mechanistic advances in axon pathfinding. Curr. Opin. Cell Biol. 63, 11–19 (2020).

57. Fricker, M., Tolkovsky, A. M., Borutaite, V., Coleman, M. & Brown, G. C. Neuronal Cell Death. Physiol. Rev. 98, 813–880 (2018).

58. Gordon, N. Apoptosis (programmed cell death) and other reasons for elimination of neurons and axons. Brain Dev. 17, 73–77 (1995).

59. Fame, R. M., MacDonald, J. L. & Macklis, J. D. Development, specification, and diversity of callosal projection neurons. Trends Neurosci. 34, 41–50 (2011).

60. Luca, A. de et al. Distinct Modes of Neuritic Growth in Purkinje Neurons at Different Developmental Stages: Axonal Morphogenesis and Cellular Regulatory Mechanisms. PLoS ONE 4, e6848 (2009).

61. Uesaka, N., Hirai, S., Maruyama, T., Ruthazer, E. S. & Yamamoto, N. Activity Dependence of Cortical Axon Branch Formation: A Morphological and Electrophysiological Study Using Organotypic Slice Cultures. J. Neurosci. 25, 1–9 (2005).

62. Szebenyi, G., Callaway, J. L., Dent, E. W. & Kalil, K. Interstitial Branches Develop from Active Regions of the Axon Demarcated by the Primary Growth Cone during Pausing Behaviors. J. Neurosci. 18, 7930–7940 (1998).

63. Bhide, P. & Frost, D. Stages of growth of hamster retinofugal axons: implications for developing axonal pathways with multiple targets. J. Neurosci. 11, 485–504 (1991).

64. Leibold, N. S. et al. NMDA receptor activation drives early synapse formation in vivo. bioRxiv 2024.05.23.595343 (2024) doi:10.1101/2024.05.23.595343.

65. Rosenberg, S. S. & Spitzer, N. C. Calcium Signaling in Neuronal Development. Cold Spring Harb. Perspect. Biol. 3, a004259 (2011).

66. McKinney, A. A., Petrova, R. & Panagiotakos, G. Calcium and activity-dependent signaling in the developing cerebral cortex. Dev. (Camb., Engl.) 149, dev198853 (2022).

67. Gasperini, R. J. et al. How does calcium interact with the cytoskeleton to regulate growth cone motility during axon pathfinding? Mol. Cell. Neurosci. 84, 29–35 (2017).

68. Andreae, L. C. & Burrone, J. The role of neuronal activity and transmitter release on synapse formation. Curr. Opin. Neurobiol. 27, 47–52 (2014).

69. Hutchins, B. I. & Kalil, K. Differential Outgrowth of Axons and their Branches Is Regulated by Localized Calcium Transients. J. Neurosci. 28, 143–153 (2008).

70. Wen, Z., Guirland, C., Ming, G. & Zheng, J. Q. A CaMKII/Calcineurin Switch Controls the Direction of Ca2+-Dependent Growth Cone Guidance. Neuron 43, 835–846 (2004).

71. Tang, F., Dent, E. W. & Kalil, K. Spontaneous Calcium Transients in Developing Cortical Neurons Regulate Axon Outgrowth. J. Neurosci. 23, 927–936 (2003).

72. Korkotian, E. & Segal, M. Release of calcium from stores alters the morphology of dendritic spines in cultured hippocampal neurons. Proc. Natl. Acad. Sci. 96, 12068–12072 (1999).

73. Gomez, T. M. & Spitzer, N. C. In vivo regulation of axon extension and pathfinding by growth-cone calcium transients. Nature 397, 350–355 (1999).

74. Cao, J.-W., Liu, L.-Y. & Yu, Y.-C. Gap junctions regulate the development of neural circuits in the neocortex. Curr. Opin. Neurobiol. 81, 102735 (2023).

75. Luhmann, H. J. et al. Spontaneous Neuronal Activity in Developing Neocortical Networks: From Single Cells to Large-Scale Interactions. Front. Neural Circuits 10, 40 (2016).

76. Blankenship, A. G. & Feller, M. B. Mechanisms underlying spontaneous patterned activity in developing neural circuits. Nat. Rev. Neurosci. 11, 18–29 (2010).

77. Moore, S. J. & Murphy, G. G. The role of L-type calcium channels in neuronal excitability and aging. Neurobiol. Learn. Mem. 173, 107230 (2020).

78. Correa, B. H. M., Moreira, C. R., Hildebrand, M. E. & Vieira, L. B. The Role of Voltage-Gated Calcium Channels in Basal Ganglia Neurodegenerative Disorders. Curr. Neuropharmacol. 21, 183–201 (2023).

79. Lepski, G., Jannes, C. E., Nikkhah, G. & Bischofberger, J. cAMP promotes the differentiation of neural progenitor cells in vitro via modulation of voltage-gated calcium channels. Front. Cell. Neurosci. 7, 155 (2013).

80. Lee, D. Global and local missions of cAMP signaling in neural plasticity, learning, and memory. Front. Pharmacol. 6, 161 (2015).

81. Hall, D. D. et al. Critical Role of cAMP-Dependent Protein Kinase Anchoring to the L-Type Calcium Channel Cav1.2 via A-Kinase Anchor Protein 150 in Neurons †. Biochemistry 46, 1635–1646 (2007).

82. Alfadil, E. & Bradke, F. Moving through the crowd. Where are we at understanding physiological axon growth? Semin Cell Dev Biol (2022) doi:10.1016/j.semcdb.2022.07.001.

83. Severino, F. P. U. et al. The role of dimensionality in neuronal network dynamics. Sci. Rep. 6, 29640 (2016).

84. Sato, M., Lopez-Mascaraque, L., Heffnerai, C. D. & O’Leary, D. D. M. Action of a diffusible target-derived chemoattractant on cortical axon branch induction and directed growth. Neuron 13, 791–803 (1994).

85. Baier, H. & Bonhoeffer, F. Axon Guidance by Gradients of a Target-Derived Component. Science 255, 472–475 (1992).

86. Molnár, Z. & Blakemore, C. Lack of regional specificity for connections formed between thalamus and cortex in coculture. Nature 351, 475–477 (1991).

87. Diaz-Cuadros, M. & Pourquié, O. The Clockwork Embryo: Mechanisms Regulating Developmental Rate. Annu. Rev. Genet. 57, 117–134 (2023).

88. Kim, J. et al. Human assembloids reveal the consequences of CACNA1G gene variants in the thalamocortical pathway. Neuron (2024) doi:10.1016/j.neuron.2024.09.020.

89. Yu, C. R. et al. Spontaneous Neural Activity Is Required for the Establishment and Maintenance of the Olfactory Sensory Map. Neuron 42, 553–566 (2004).

90. Kandel, E. R. The molecular biology of memory: cAMP, PKA, CRE, CREB-1, CREB-2, and CPEB. Mol. Brain 5, 14 (2012).

91. Barbado, M., Fablet, K., Ronjat, M. & Waard, M. D. Gene regulation by voltage-dependent calcium channels. Biochim. Biophys. Acta (BBA) - Mol. Cell Res. 1793, 1096–1104 (2009).

92. Munno, D. W., Prince, D. J. & Syed, N. I. Synapse Number and Synaptic Efficacy Are Regulated by Presynaptic cAMP and Protein Kinase A. J. Neurosci. 23, 4146–4155 (2003).

93. West, A. E. et al. Calcium regulation of neuronal gene expression. Proc. Natl. Acad. Sci. 98, 11024–11031 (2001).

94. Klocke, B. et al. Insights into the role of intracellular calcium signaling in the neurobiology of neurodevelopmental disorders. Front. Neurosci. 17, 1093099 (2023).

95. Pourtavakoli, A. & Ghafouri-Fard, S. Calcium signaling in neurodevelopment and pathophysiology of autism spectrum disorders. Mol. Biol. Rep. 49, 10811–10823 (2022).

96. Delhaye, S. & Bardoni, B. Role of phosphodiesterases in the pathophysiology of neurodevelopmental disorders. Mol. Psychiatry 26, 4570–4582 (2021).

97. Lorenzon, N. M. & Beam, K. G. Calcium channelopathies. Kidney Int. 57, 794–802 (2000).

98. Ferron, L. & Zamponi, G. W. The road to the brain in Timothy syndrome is paved with enhanced CaV1.2 activation gating. J. Gen. Physiol. 154, e202213272 (2022).

99. Paşca, S. P. et al. Using iPSC-derived neurons to uncover cellular phenotypes associated with Timothy syndrome. Nat. Med. 17, 1657–1662 (2011).

100. Brimson, C. A. et al. Collective oscillatory signaling in Dictyosteliumdiscoideum acts as a developmental timer initiated by weak coupling of a noisy pulsatile signal. Dev. cell (2024) doi:10.1016/j.devcel.2024.11.016.

101. Dana, H. et al. High-performance calcium sensors for imaging activity in neuronal populations and microcompartments. Nat. Methods 16, 649–657 (2019).

102. Yau, K. W. et al. Dendrites In Vitro and In Vivo Contain Microtubules of Opposite Polarity and Axon Formation Correlates with Uniform Plus-End-Out Microtubule Orientation. J. Neurosci. 36, 1071–1085 (2016).

103. Sutcliffe, M. & Lancaster, M. A. A Simple Method of Generating 3D Brain Organoids Using Standard Laboratory Equipment. Methods Mol. Biol. (Clifton, NJ) 1576, 1–12 (2017).

104. Qin, D., Xia, Y. & Whitesides, G. M. Soft lithography for micro- and nanoscale patterning. Nat. Protoc. 5, 491–502 (2010).

105. Brittain, S., Paul, K., Zhao, X.-M. & Whitesides, G. Soft lithography and microfabrication. Phys. World 11, 31–37 (1998).

106. Fujii, T. PDMS-based microfluidic devices for biomedical applications. Microelectron. Eng. 61–62, 907–914 (2002).

107. Duffy, D. C., McDonald, J. C., Schueller, O. J. A. & Whitesides, G. M. Rapid Prototyping of Microfluidic Systems in Poly(dimethylsiloxane). Anal. Chem. 70, 4974–4984 (1998).

108. Preibisch, S., Saalfeld, S. & Tomancak, P. Globally optimal stitching of tiled 3D microscopic image acquisitions. Bioinformatics 25, 1463–1465 (2009).

109. Wolf, F. A., Angerer, P. & Theis, F. J. SCANPY: large-scale single-cell gene expression data analysis. Genome Biol. 19, 15 (2018).

110. Setty, M. et al. Characterization of cell fate probabilities in single-cell data with Palantir. Nat. Biotechnol. 37, 451–460 (2019).

111. Sumanaweera, D. et al. Gene-level alignment of single-cell trajectories. Nat. Methods 22, 68–81 (2025).

112. Prigge, C. L. & Kay, J. N. Dendrite morphogenesis from birth to adulthood. Curr. Opin. Neurobiol. 53, 139–145 (2018).

113. Felipe, J. D., Marco, P., Fairén, A. & Jones, E. G. Inhibitory synaptogenesis in mouse somatosensory cortex. Cereb. cortex (N. York, NY : 1991) 7, 619–634 (1997).

